# Coupled SDE-ODE Modeling of Tumor-Immune Dynamics to Infer Biomarker Release

**DOI:** 10.1101/2025.08.15.670571

**Authors:** Pujan Shrestha, Yijia Fan, Jason T. George

## Abstract

Tumor–immune interactions are central to cancer progression and treatment response, driving cell death through immune-mediated killing and resource-limited competition. In early-stage disease or following effective treatment, cancer populations are often small and difficult to observe directly. Disease monitoring therefore relies on the detection of biomarkers such as circulating tumor DNA (ctDNA) as noisy proxies to cancer size. However, existing approaches lack robust frameworks to infer tumor burden from these signals when populations approach detection thresholds. To address this, we present a coupled deterministic–stochastic framework that links tumor–immune dynamics to biomarker release. A two-prey, one-predator Lotka–Volterra model captures interactions between immune cells and competing tumor subpopulation under shared resource constraints. Biomarker production is modeled using a linear stochastic differential equation, incorporating ctDNA release from both apoptotic death (immune-mediated) and necrotic death (due to intra-tumor competition). We derive analytical solutions for the resulting biomarker trajectories, including mean detection time under a minimal detection threshold. These results show how volatility in measured biomarker signal, disease heterogeneity, and immune pressure jointly shape signal emergence and persistence. Finally, to model situations in which the observer has access to future information—such as the terminal biomarker signal or sampling time in retrospective studies—we adopt an anticipative theoretic perspective. Using anticipative stochastic calculus, we derive path solutions to the resulting anticipating stochastic differential equation, capturing how future observations influence the inferred biomarker dynamics. This approach links the dynamics of underlying tumor–immune interactions to the corresponding detectable biomarker levels, with implications for early detection, immune monitoring, and retrospective reconstruction of disease progression.

---

The adaptive immune system plays an important role in shaping cancer development, and tumor-immune interactions lead to one of three possible clinical outcomes: tumor escape, tumor elimination, or a period of sustained equilibrium (1, 2). In the elimination phase, CD8+ cytotoxic T lymphocytes (CTL) and natural killer (NK) cells recognize and destroy tumor cells by recognizing tumor-associated antigens (TAA) presented on tumor major histocompatibility complex (MHC) (1). Cancer cells may enter into a state of equilibrium with an active immune system if they can avoid complete destruction. Such states are reflected by interesting dynamical properties that give rise to potentially large waiting times (3, 4). Over time, malignant cells can become more immune evasive through a variety of mechanisms, including down-regulation of antigen presentation machinery, up-regulation of immune checkpoint molecules (e.g. PD-L1), loss of recognized tumor-associated antigens, and secretion of immuno- suppressive cytokines (2, 5–9).

Despite growing insights into tumor–immune interactions, a current major limitation to understanding these details more precisely remains in the ability to accurately infer tumor burden from circulating biomarkers, particularly when disease levels are low (10). One existing and clinically relevant example is circulating tumor DNA (ctDNA) released during apoptotic and necrotic cell death. However, its interpretation is complicated by stochastic shedding dynamics, contributions from heterogeneous tumor subpopulation, and noise introduced by sampling and clearance processes (11– 13). These factors become especially limiting near detection thresholds, where signal-to-noise ratios deteriorate and conventional inference methods often fail to recover underlying tumor dynamics (14, 15). A key unmet need is a principled framework that links observed biomarker fluctuations to population-level tumor behavior, particularly under conditions of uncertainty. Analogous challenges arise in chronic infections such as HIV and hepatitis C, where low-level residual disease may persist below detection yet remain biologically relevant (16, 17).

To address these challenges, we develop a hybrid mathematical framework that couples deterministic tumor–immune dynamics with an underlying stochastic model of biomarker release. We introduce an ordinary differential equation (ODE) model of tumor–immune interaction based on a predator–prey (immune-tumor) structure (18–23). Cancer cells are divided into immune-targeted and immune-evasive subpopulations, allowing the model to capture selection under variable fitness costs incurred under immune pressure. Immune-targeted cells are eliminated through immune-mediated apoptosis, represented by Lotka–Volterra-type interaction terms, while logistic growth constraints lead to necrotic death under high tumor burden (24, 25). Building on this foundation, we introduce a stochastic differential equation (SDE) that links tumor cell death—via both apoptosis and necrosis—to the generation of ctDNA and other measurable biomarkers (26–29). This stochastic component captures the intrinsic variability in biomarker shedding, clearance, and measurement noise. Apoptotic and necrotic processes contribute to ctDNA through distinct mechanisms (for example, immunemediated fragmentation versus passive lysis) allowing the model to distinguish their molecular signatures.

By modeling heterogeneity in immune targeting under shared resource constraints, we capture a range of escape and coexistence outcomes that depend on mutual altruism and egoism between the subpopulations under differential immune targeting. We then extend this system using stochastic differential equations to track the noisy evolution of biomarker signals, including those arising from apoptosis and necrosis. This formulation allows us to analytically characterize when and how such signals become detectable, and to identify key parameters, including volatility, that influence early detection. Finally, we introduce an anticipative stochastic integration approach that incorporates future data to obtain biomarker trajectories, offering a complementary perspective on retrospective inference. Together, our findings provide a theoretical foundation for interpreting noisy biomarker data in the context of immune surveillance, and suggest design principles for improving detection of minimal residual disease.

## Methods and Materials

### Model Development

We present a generalized framework for modeling interactions between a heterogeneous population of cancer cells and the immune system, combining deterministic tumor growth and immune response dynamics with stochastic variability in biomarker release. We assume that the tumor subpopulations and T cells behave according to a one-predator, two-prey Lokta-Volterra model. Furthermore, we assume that, in addition to T cell elimination pressure, the two tumor subpopulations are also competing with each other under a shared carrying capacity with parameters denoting the degree of altruism vs egoism. Such social behavior like altruism has been previously documented in breast cancers and is a subject of active research (30). In a previous work, the authors constructed a mathematical model where immunomodulation would impact the per-cell killing rate of the birth-death process representing tumor population (31). Under this context, this immunomodulation can be represented as changes in the T cell interaction and expansion terms.

#### Tumor Immune Dynamics

Let *B, E*, and *I* represent the baseline tumor, evasive tumor, and T cells compartments, respectively. The three compartments are linked via a system of ODE’s, given by:

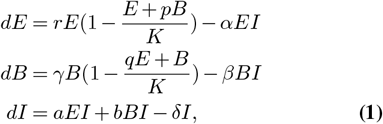

with *E*(0),*B*(0),*I*(0) > 0, and r, *γ, α, β*, a, b, *δ* > 0 as the model parameters. We consider a finite time horizon as such, 0 ≤ *t* ≤ *T* for *T ≤* 0. The parameters definitions are given in Table 1. We define the competitive parameters *p* and *q* to represent the relative level of altruism/egoism of each subpopulation, as has been modeled previously (32). In this interpretation, egoistic and altruistic behaviors could be seen as as regimes for the parameters. For example, when *q* > 1, the baseline tumor puts more weight on the evasive tumor volume in modulating its own growth rate. On the other hand, 0 < *q* < 1 would induce the opposite behavior.

**Table 1.** Model parameters and their descriptions

| Parameter(s) | Description |
| --- | --- |
| $K$ | Carrying capacity for the tumors |
| $r, \gamma$ | Tumor growth rates |
| $\alpha, \beta$ | Tumor-immune interaction tumor death rates, with $\alpha < \beta$ |
| $a, b$ | T cell growth rates resulting from tumor-immune interaction |
| $\delta$ | T cell death rate |
| $p, q$ | Scaling parameters for competition between cancer populations under shared carrying capacity |

We remark that the high number of parameters presents a challenge for future data fitting and modeling. To mitigate this, we perform time scaling and population scaling relative to a characteristic immune compartment size *I*0 to obtain an alternate dimensionless version of the model with a reduced number of (six) parameters, given by Eq. (1). For more details, refer to Sec S4.3 of SI.

#### Biomarker Dynamics

The detailed physiological processes underlying biomarker kinetics is complex and often involves multiple independent contributions to overall dynamics. For foundational understanding, we motivate our framework by considering the example of ctDNA - a biomarker which has previously been used as a specific indicator of tumor burden and is often most informative when measured in the regime of low tumor cell populations, near the detection limit. Moreover, ctDNA release into the bloodstream occurs through apoptosis or necrosis, with the release rate being approximately proportional to tumor size (26, 27, 33, 34). Similarly, (primary renal and hepatic) clearance mechanisms follow first-order kinetics, implying that the rate of biomarker decay is proportional to its current concentration (28, 35). Noise in these systems arising from micro-environmental fluctuations, immune interactions, or patient-specific factors is also proportional to the biomarker level, as suggested in prior stochastic modeling frameworks of ctDNA dynamics (36). Consequently, under the assumption that biomarker levels have log-normal distribution and multiplicative behavior, we use a family of linear SDEs to capture both deterministic influences from tumor-immune interactions and inherent biological variability in biomarker levels and their consequent detection. Namely, *C*(*t*) tracks biomarker shedding levels from the tumor-immune dynamics given in Eq. (1). *C*(*t*) may be represented by the SDE:

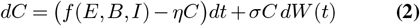

where *C*(0) > 0, *W* (*t*) is standard Brownian motion and *σ η* are the volatility and the decay rates, respectively. We assume that apoptotic events occur via the interactions between tumor and T cells. As such, we consider such interactions in the appropriate tumor subpopulation as well as the joint release for the shedding mechanism. Similarly, the necrotic release is tracked by the “over-crowding” term that arises from the shared carrying capacity. Using the longitudinal population dynamics from the ODE model, we model the apoptotic and necrotic release for each tumor subpopulation. Consequently, the total biomarker release summed over evasive/baseline and apoptotic/necrotic contributions becomes a linear combinations of these interaction terms. In Table 2, we list different functional forms of interest for *f*. Note that, the model assumes a universal decay and volatility factor common to all release mechanisms. For this paper, we assume that *W* (*t*) in the SDE for one release mechanism is independent and identically distributed to the Brownian noise of the other SDE’s.

**Table 2.** Biomarker trajectory expressions for different tumor and cell death dynamics.

| Biomarker | Expression $f(E, B, I)$ |
| --- | --- |
| Evasive Tumor | $rE \left( \frac{E+pB}{K} \right) + \alpha EI$ |
| Baseline Tumor | $\gamma B \left( \frac{qE+B}{K} \right) + \beta BI$ |
| Apoptotic Cells | $\alpha EI + \beta BI$ |
| Necrotic Cells | $\gamma B \left( \frac{qE+B}{K} \right) + rE \left( \frac{E+pB}{K} \right)$ |

### Numerical Simulations

The ODE was solved and generated via the numpy and scipy library on Python. For the stochastic differential equations, the numerical so-lution simulations were performed via Euler-Maruyama scheme and path solution simulations were generated via Monte Carlo simulations. Github code for the model is available at https://github.com/pujantamu/ODE_SDE_Model

### Data Analysis

To extend our investigation beyond the predictions of the ODE-SDE model, we implemented a two-dimensional agent-based model (ABM) to simulate stochastic tumor-immune interactions using the Gillespie stochas-tic simulation algorithm (37). This approach enables us to test how cell-level stochasticity in immune efficacy can drive divergent outcomes. In this model, individual tumor cells and cytotoxic T cells are represented as discrete agents, each with prespecified behavior. Tumor cell behavior includes two-dimensional random migration and proliferation, while T cell behavior includes chemotaxis-driven migration toward regions of high chemokine concentration secreted by tumor cells, as well as tumor cell killing and clonal expansion (38). T cell mediated killing occurs when a T cell is within a defined interaction radius of a tumor cell, representing physical proximity required for cytotoxic engagement (39). Once this spatial condition is met, tumor cell killing proceeds probabilistically, governed by a predefined T cell killing probability. Upon successful killing, the T cell undergoes clonal expansion. The model assumes a homogeneous tumor cell population without phenotypic and mutational variability. The rates of each event were determined based on biological plausibility (40, 41). This approach enables the modeling of celllevel stochasticity and discrete interaction events, offering a mechanistic representation of tumor-immune dynamics beyond the deterministic framework.

## A. Results

Below, we summarize our key findings. We refer the reader to the SI for complete mathematical details.

### A.1. Dynamical Behavior of the Tumor-Immune System

*Theorem 1* (Existence and Uniqueness) The solution of the system of ODE given by Eq. (1) exists, is unique, and depends on the initial conditions. Furthermore, if the initial conditions are positive, then solutions are restricted to the positive octant.

*Theorem 2* (Equilibrium states and Local stability) The equilibrium states alongside their existence and stability conditions for the system of ODE given by Eq. (1) are given in Table 3. Here *p*0, *q*0, *pmax, qmax* are constants dependent on the parameter values. Furthermore, for ⋆, we have to apply the Routh-Hurwitz stability criterion (42) on the characteristic equation for the interior state to obtain stability conditions.

**Table 3.** Parameter dependence on existence and stability of Equilibrium states

|  | Equilibrium States | Existence | Stability |
| --- | --- | --- | --- |
| (i) | $(0, 0, 0)$ | Exists | Unstable |
| (ii) | $(K, 0, 0)$ | Exists | $q > 1$ and $K < \frac{\delta}{a}$ |
| (iii) | $(0, K, 0)$ | Exists | $p > 1$ and $K < \frac{\delta}{b}$ |
| (iv) | $\left(\frac{K(1-p)}{1-pq}, \frac{K(1-q)}{1-pq}, 0\right)$ | $pq \neq 1$ and<br>$(p, q > 1 \text{ or } p, q < 1)$ | $\delta > \frac{K}{1-pq} (a(1-p) + b(1-q))$ |
| (v) | $\left(\frac{\delta}{a}, 0, \frac{r}{\alpha} \left(1 - \frac{\delta}{aK}\right)\right)$ | $K > \frac{\delta}{a}$ | $q > q_0$ |
| (vi) | $\left(0, \frac{\delta}{b}, \frac{\gamma}{\beta} \left(1 - \frac{\delta}{bK}\right)\right)$ | $K > \frac{\delta}{b}$ | $p > p_0$ |
| (vii) | $(\hat{E}, \hat{B}, \hat{I})^*$ | $(p, q > p_0, q_0 \text{ or } p, q < p_0, q_0)$<br>and $p, q < p_{max}, q_{max}^*$ | ★ |

*Proof:* Section 4.1 & 4.2 of the SI.

From Theorem 1, restriction of the solution to the positive octant allows us to find parametric restrictions for existence of the equilibrium points. Theorem 2 provides an analytical perspective on the local stability of these equilibrium points. We describe these states below, which are summarized in Table 3.

The state where all compartments vanish (Table 3(*i*)) is unstable for all parametric combinations and thus is unreachable in the deterministic system, provided that there is a nonzero starting population in each compartment. In contrast, the tumor competition parameters play a key role in the existence and stability of the remaining equilibrium states. When looking at evasive homogeneous escape state in Table 3(*ii*), the immune evasive cancer population grows to carrying capacity with mutual extinction of both baseline cancer and adaptive immune populations whenever a) the baseline tumor exhibits altruism (*q* > 1) and b) the ratio of the T cell death rate to the T cell growth rate via the interactions between the T cells and the evasive tumor exceeds the carrying capacity. Interchanging the roles of parameters specific to the baseline and evasive cancer subpopulation give rise to a solitary baseline cancer population growing to carrying capacity (Table 3(*iii*)).

In heterogeneous tumor escape state, Table 3(*iv*), see the role of the competitive parameters play shaping this state. Both the baseline and evasive equilibrium states themselves are scaled by the competition parameters, while existence requires both tumor populations would have to be simultaneously either egoistic (*p*< 1, *q* < 1) or altruistic (*p*> 1, *q* > 1) for such a condition to even exist. This result aligns well with the game-theoretic intuition that populations with mixed strategies would lead to the dominance of the egoistic strategist.

The homogeneous tumor-immune coexistence states (Table 3(*v*),(*vi*)), wherein one of the tumor subpopulation strikes a balance with the immune system while the other tumor population faces extinction, provide additional insight into the interplay of the competitive forces of the tumor subtypes. We obtain a lower bound on the competition parameters, *p*0 or *q*0, required for stability in the cases where only one cancer subpopulation survives. In particular, the dying subpopulation must exhibit altruism in excess of a lower bound on the population’s respective competition parameter. These lower bounds on can be obtained from SI Eqs. S13 and Eq. S14 and are given by

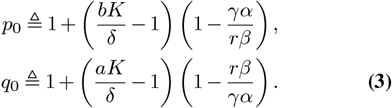

These bounds are modulated by the relative competitive advantage between the cancer subpopulations and the relative free space left under shared carrying capacity. Moreover, they act independently for the homogeneous tumor-immune coexistence state. However, we find a mutual coaction being required for the interior equilibrium state given by Table 3(*vii*). This is perhaps the most interesting state, wherein both cancer subpopulations co-exist with the immune compartment under a shared carrying capacity leading to a heterogeneous immune-mediated coexistence state. Existence requires both tumor populations to be simultaneously either more egoistic or more altruistic than the lower bounds given in Eq. (3). More concretely, if *p > p*0, *q* > *q*0 and the evasive cancer subpopulation enjoys a competitive advantage (*β* > *α, r* > *γ*), then the evasive tumor is more likely to fully occupy the carrying capacity leading to a homogeneous coexistence state or immune escape state. We can use the relation *r β* > *γ α* to obtain *p*0 > 1 > *q*0. Thus, the evasive tumor has to be altruistic (*p* > *p*0 > 1) while the baseline tumor is allowed a little bit of egoism (*q* ∈ (*q*0, 1) ∪ [1, ∞)). The interior state therefore requires a balance between the immune forces and the competitive forces for existence and stability. Note that, the tumor-immune coexistence state when considering low tumor populations overlaps with empirically observed immune-mediated dormancy (4).

Intriguingly, *pq* ≠ 1 emerges as a critical condition for immune-mediated dormant states for which the tumor subpopulations strike a balance with the immune compartment resulting in a net zero growth rate. Indeed, for the interior equilibrium with non-zero values, *p* = *q* = 1 would imply that either, at equilibrium, *I* = 0 contradicting the non zero assumption or there exists a constant *k*0 such that *γ α* = *k*0*r β*. In the latter case, *dE* = *k*0*dS* which would mean that the evasive population is nothing but a scaled version of the baseline one. Under this very restrictive condition on the competitive rates, the two cancer subpopulations match exactly in their dynamics and as such, can be viewed as a single (homogeneous) population.

### A.2. Analysis and comparison with agent-based modelling

We analyze the baseline (immune-sensitive) cancer compartment as a representative example. Specifically, we investigate how variations in tumor-immune interaction parameters (*α* and *a*) affect system equilibria. With low interaction and T cell expansion, the tumor escape outcome, (0,*K*, 0), exists stably. As the value of the interaction parameter *α* increases, we find that the coexistence equilibrium becomes reachable, and its stability then depends on the competitive parameters given in Eq. 3. As *α* ranges from zero to one, we have that *p*0 is a decreasing function of *α* if 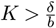 and increasing otherwise. Namely,

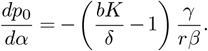

Thus, the interactions governed by *α* and *a* are key determinants of stable equilibrium states, and changes to these values allow for cancer immune escape, equilibrium, and possible elimination (1).

To examine how stochastic immune behavior at the cellular level influences tumor dynamics, we simulate the tumor-immune interaction using a previously developed agent-based model (ABM) tracing the tumor-immune interaction (37). In the ABM, the tumor-immune interaction parameters correspond to the initial number of T cells and the predefined T cell killing probability. We explored the impact of varying these two parameters while fixing all other factors. Simulations were conducted across a 10 11 grid representing different combinations of initial T cell counts and T cell killing probabilities. For each parameter combination, the ABM was executed and the tumor burden was recorded after a fixed simulation time (*t* = 100). The resulting heatmap (Fig 2) shows the average tumor burden at the end of the simulation, with color intensity representing tumor burden magnitude. We defined outcome regions based on the final tumor burden: 1) Elimination (dark blue): tumor burden approaches zero; 2) Equilibrium (orange): intermediate, stabilized tumor burden; 3) Escape (red): tumors attain 90% of their carrying capacity. Representative time-course plots for selected parameter combinations for each category are shown in panels B-I, illustrating dynamic tumor burden and T cell population changes over time for different regimes. Notably, tumor elimination is only observed in regions with both sufficiently high initial T cell counts and high killing probability, suggesting a synergistic requirement for both T cell abundance and cytotoxic efficiency in achieving immune-mediated tumor clearance. In contrast, low initial T cell numbers or low killing probabilities consistently lead to tumor escape, indicating that deficits in either recruitment or function of T cells can independently compromise tumor control. The intermediate region corresponds to a dynamic equilibrium, where immune pressure slows tumor growth but is insufficient for complete elimination. This mirrors clinically observed tumor-immune steady states and may represent a window of opportunity for combination therapies to tip the balance toward elimination. The sharp transitions between outcome regions also highlight the system’s nonlinear sensitivity to immune parameters, implying that small changes in immune cell behavior due to, for instance, immunotherapy or tumor-induced suppression, can result in dramatically different clinical outcomes.

**Fig. 1.**
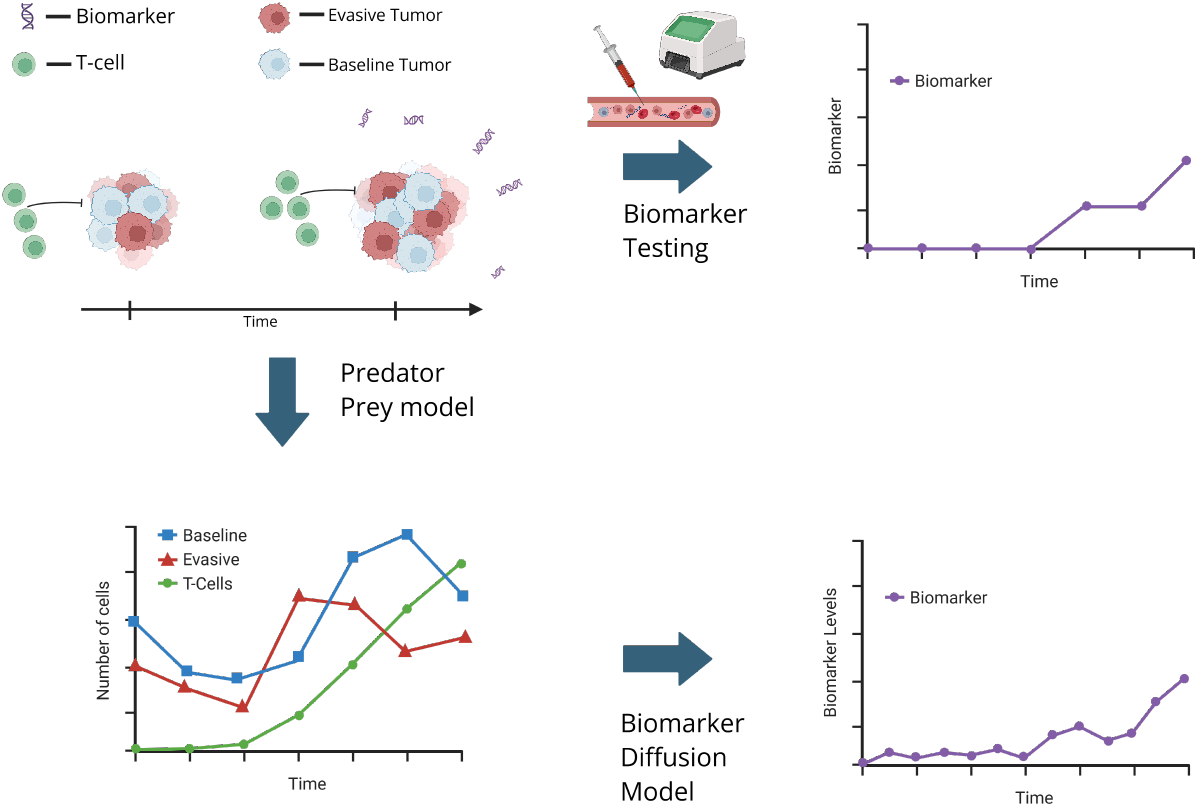
Conceptual schematic for the ODE-SDE model in biomarker trajectories. Tumor populations evolve under immune pressure, giving rise to heterogeneous dynamics between an immune-sensitive ‘baseline’ subpopulation and immune-evasive cells. These interactions, along with immune-mediated killing, drive biomarker release into circulation through apoptotic and necrotic cell death. While the underlying population dynamics (bottom left) remain unobservable, they influence measurable biomarker levels (top right), which exhibit stochastic fluctuations due to variability in shedding and clearance. Coupling deterministic tumor-immune dynamics with stochastic biomarker shedding provides a possible framework for inferring hidden cancer population states from noisy biomarker data.

**Fig. 2.**
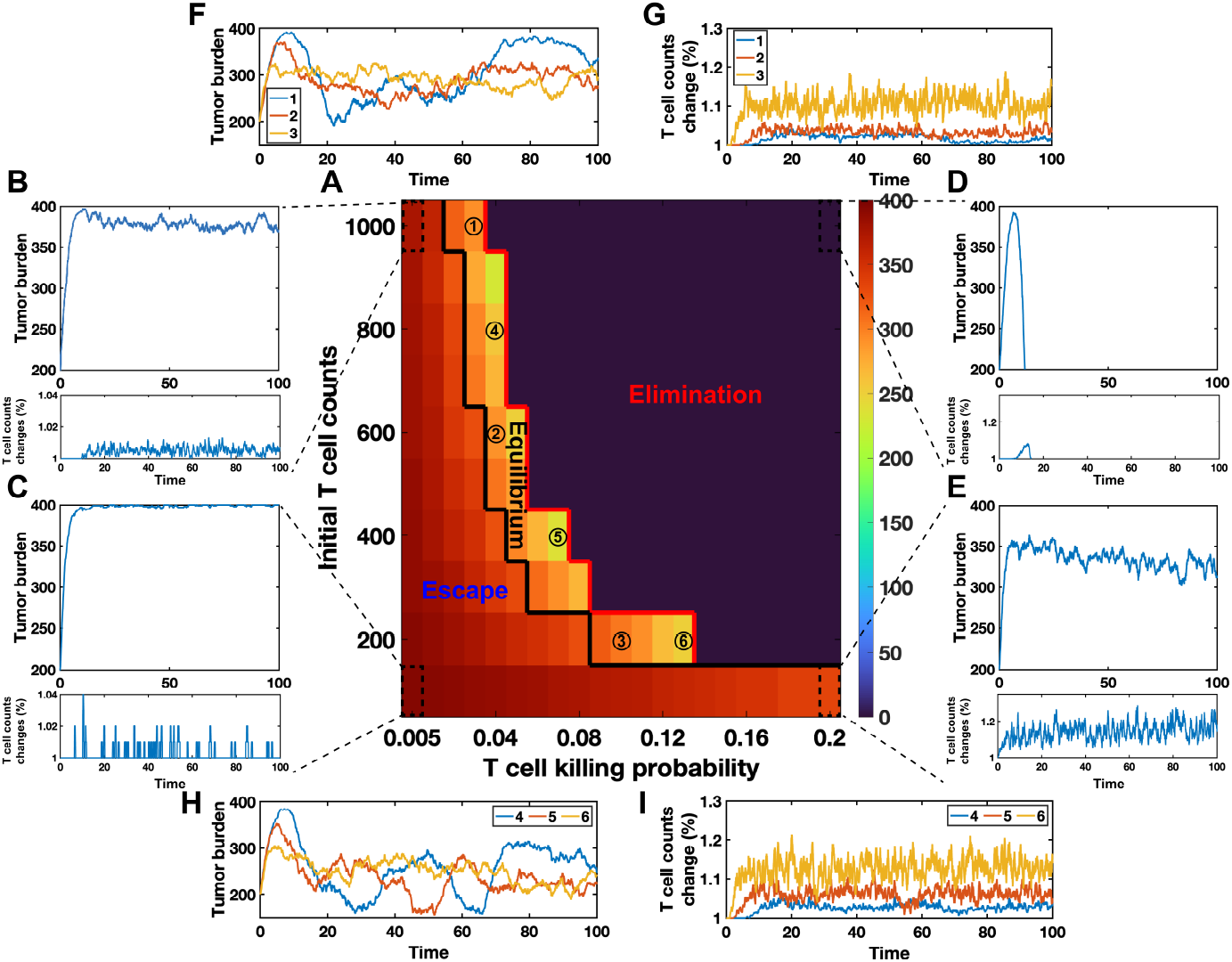
Heatmap showing tumor burden outcomes across a range of immune interaction parameter regimes in ABM. A) 10 × 11 parameter grid was explored by varying the initial T cell counts (y-axis) and predefined T cell killing probability (x-axis), while all other model parameters were held constant. The color scale represents the average tumor burden at simulation time t = 100, with dark colors indicating lower tumor burden. Three distinct outcome regions are observed: Elimination (dark blue), equilibrium (orange), and escape (red), separated by black and red contour lines. B)-I): Representative time-course dynamics of tumor burden and T cell counts for selected parameter regimes.

While the parameters are not one-to-one for the ODE model with respect to the ABM, we postulate that some of the parameters can be inferred as well as related. For example, the expansion rate for the tumor in the ABM can infer the growth rate of the tumor compartment. Here, we assume that T cell killing probabilities represent interaction and expansion parameters. As the T cell probabilities change, we are able to observe an analogous set of outcomes as a way to investigate our results derived from theory.

### A.3. Global Stability of the Tumor-Immune System

The system of ODEs given in Eq. (1) is nonlinear and as such, stability analysis via the linearized system is restricted to describing the behavior of trajectories close to equilibrium. We investigate global stability for the interior equilibrium state via Lyapunov’s second method of stability.

*Theorem 3* (Lyapunov Stability) Consider the following functional *L* given by

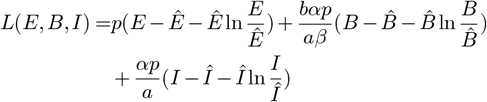

where 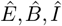 are the interior state values. Then *L* is positive definite and zero only at the interior equilibrium. Furthermore, 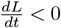 and consequently, *L* is a Lyapunov functional when either of the following conditions hold:

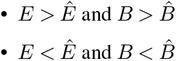

*Proof:* See Sec S4.2 of SI

The functional form of *L* was chosen to measure deviations from the equilibrium while allowing for easier tractability with the time derivative operator. We find that trajectories where the tumor population size is either jointly above or below the interior equilibrium size will stability converge to the equilibrium globally asymptotically. However, fluctuations in the trajectories when the above conditions are not satisfied might lead to possible extinction of at least one subcompartment.

### A.4. Dynamical Behavior of the Biomarker Compartment

So far we have carefully analyzed the deterministic arm of the model which is meant to track tumor-immune interactions. To get a better understanding of how the biomarker release reflects the longitudinal tumor-immune dynamics, we consider the stochastic differential equation given by Eq. (2) and assume that the drift terms for the biomarker release mechanisms are driven by apoptotic and necrotic events in the tumor immune microenvironment: apoptotic via tumor-immune killing, necrotic via carrying capacity-induced cell death. Let *c*^*^ be the minimal biomarker size required for the tumor to be detected. The model dynamics from the competitive Lokta-Volterra system serve as an input to the time-dependent drift for the biomarker model. We remark that the general form of the drift term enables the modeling of the biomarker trajectory in multiple biological contexts depending on the nature of the signal source. For foundational understanding, we normalize the biomarker level by *c*^*^. Under this assumption, we consider the possible forms and rationale for the drift term *f* (Table 4). Note that this is the same release assumptions albeit normalized as in Table 2: apoptotic via tumor-immune interaction and necrotic via carrying capacity related deaths.

**Table 4.**
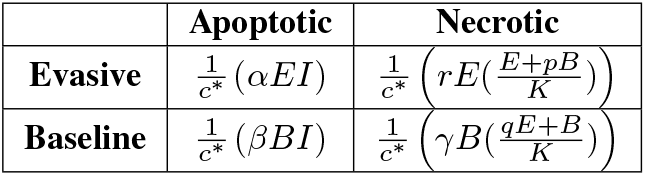
Form of *f* for Evasive and Baseline Tumors

The drift term *f* in the SDE is then a linear combination of these forms. For example, when considering the total apoptotic biomarker levels, we can sum the two functional forms for baseline and evasive apoptotic tumors and use the resulting sum as the drift. The resulting drift equation is sum of a product of Lipschitz functions and as such, it is also Lipschitz. We can use the results from Theorem 1 to obtain a unique solution (up to probability) for Eq. (2). Namely, *Theorem 4* (Existence and Uniqueness) Given that *E, B, I* are solutions of Eq. (1), the drift term *f* is a linear combination of the terms in Table 4 and consider the stochastic differential equation given by Eq. (2), then the solution exists, is unique almost surely, and given by

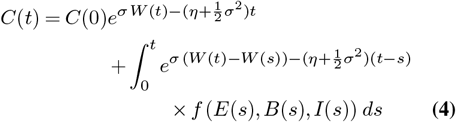

Furthermore, by the construction of *f* as sums and products of non-negative functions, *C*(*t*) is non-negative.

*Proof:* Section S5.1.1 of the SI.

*Theorem 5* (Mean and Variance) The mean and variance of the stochastic differential equation given by Eq. (2) is given by

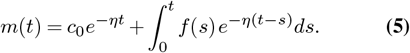

and,

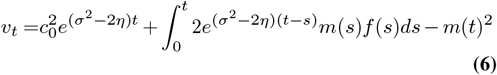

respectively.

*Proof:* Section S5.1.2 of the SI.

The above analytical characterization allows us to characterize the dynamics of biomarker release derived from the underlying tumor-immune interaction. Via Theorem 4, and Theorem 5, we generate comprehensive trajectories for all equilibrium states and list them in Section S6 of the SI. One of the key advantages of our approach is that it produces path-wise solutions for each drift condition. Given a particular realization of the stochastic noise, we obtain a complete trajectory of the biomarker dynamics, capturing fluctuations, sharp transitions, and rare events that are often obscured in expectation-based analyses. These sample paths retain the full influence of volatility on the system — including its role in shaping the variability, extremes, and temporal structure of the output. In addition, these path-wise solutions also allow for easier generalizations to the anticipative stochastic calculus.

We simulated both the path solutions obtained and the stochastic differential equation via different methods to validate the result. The SDE was numerically solved by the Euler-Maruyama scheme while the exact path solutions enables us to use Monte-Carlo simulations to generate the path trajectories. Furthermore, we use the mean and variance equations to generate the mean trajectories and the variance cone. Tumor escape states with immune loss lead to a saturation of necrotic biomarker levels as the subpopulations overcrowd the shared carrying capacity (Figures S13-S15C). On the other hand, immune-mediated coexistence generates a surplus of apoptosis-derived signal due to continued interactions between the adaptive immune and cancer compartments (Figures 3C, S11, S12C). For our purposes, we provide a full analysis in this case, which involves all three populations (Figure 3). As the tumor approaches the interior equilibrium point, the population levels fluctuate in the predator-prey dynamics (Figure 3A). These dynamics subsequently generate a cumulative biomarker release trajectory (Figure 3B), which can be further partitioned by release mechanism (Figure 3C), cancer subpopulation (Figure 3D), or both (Figure 3E). In this particular example, we observe that heterogeneous immune mediated coexistence is driven primarily via T cell killing in this visualization, which represents cases where the tumor size is small with respect to the carrying capacity. Note that *c*^*^ represents the minimal detection limit for the circulating biomarker instrument. Under the model parametrization with *c*^*^ = 100 and equilibrium state (*E, B, I*)= (2, 7, 6), the biomarker trajectory might not hit detection size at all. Consequently, we would expect such a tumor-immune interaction to provide minimal benefit to early detection, which might result in undetected cancer progression. Here, *c*^*^ was chosen arbitrarily for foundational understanding.

**Fig. 3.**
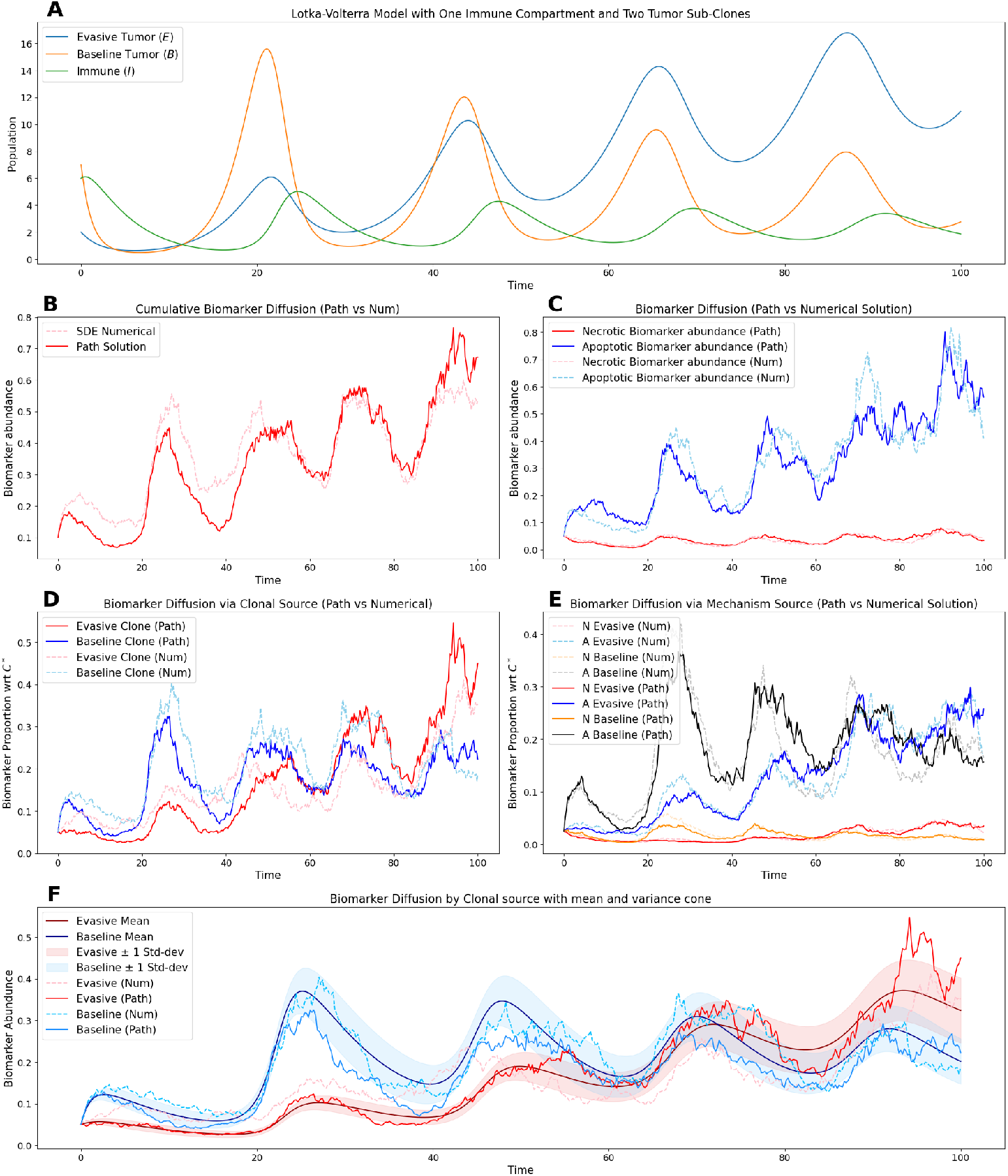
Dynamics of the heterogeneous immune mediated coexistence and associated biomarker release trajectories A) Lokta-Volterra Trajectory B) Cumulative biomarker trajectory C) Necrotic vs Apoptotic Split D) Evasive vs Baseline Split E) Full table split F) Mean and Variance of the biomarker trajectories. Dotted lines represent numerical simulations of Eq. (2) via Euler-Maruyama, lines represent path simulations of Eq. (4) via Monte Carlo simulations. In all cases, *r* = .28, *γ*= 0.5, *α*= 0.1, *β* = 0.2,a = 0.00195,*b* = 0.04, *δ* = 0.2,*p* = 0.1,*q* = 0.09,K = 100, *σ* = 0.1, *σ*= 0.1,(*E*_0_, *B*_0_, *I*_0_)= (2, 7, 6),*C*_0_ = 0.1,*C*^*^ = 100 were taken as model parameters.

Finally, we consider the model where the noise term *σCdW* (*t*) is replaced with 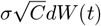. The resulting SDE is a general form of the Cox-Ingersoll-Ross (CIR) process with a positive drift 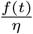 (43, 44). However, the Feller condition for the positivity of the solution is not guaranteed for such a time dependent drift. In addition, the variance grows sub-linearly when compared to the proportional growth provided by the linear version. More importantly, in transitioning from the linear variant to the non-linear one, we give up the analytical tractability of the path-wise solution. Refer to Section S6 of the SI for more details for the non-linear SDE. As such, we focus on the linear stochastic differential equation paradigm for the biomarker release.

### A.5. Impact of Volatility to time to detection

The prior section demonstrated an example for which the cancer population is expected to fall below lower detection size for a significant amount of time. Since early time plays such a critical role in cancer diagnosis and treatment monitoring, we now look into how volatility as well as the minimum detection limit impact time to detection. To accomplish this, we use the the form of the stochastic differential equation given in Eq. (2) alongside techniques from Itô theory to describe the expected time needed for the tumor to be detected under equilibrium dynamics for the LV model. We denote *f* ^*^ as the value that the drift term fixates on as the equilibrium is reached.

*Theorem 6* (Mean time to detection (*C*^*^)) Let *τ* = inf{*t* > 0 | *C*(*t*) ≥ 1} be a stopping time and let *f*(*t*) ≈ *f* ^*^. Then the time to detection is given by

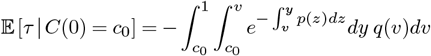

where,

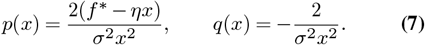

*Proof:* Section S5.2.1 of the SI.

From the form of the solution, as well as the expressions for the mean and variance of the stochastic differential equation in Theorems 4 and 5, it is evident that volatility plays a crucial role in the reachability of outcomes. Specifically, increased volatility leads to earlier readout times for the biomarker. In the regime where *σ*^2^ > 2*η*, the geometric Brownian motion structure of the SDE results in both the mean and variance increasing over time.

This behavior is evident in both simulated trajectories and in the mean time to escape, under the same underlying heterogeneous immune-mediated coexistence state studied previously. In Figure 5, we observe that higher volatility, under otherwise identical dynamics, yields a faster biomarker readout within the same time frame. Furthermore, Figure 4 illustrates the sensitivity of the mean time to escape to changes in the minimal detection threshold.

**Fig. 4.**
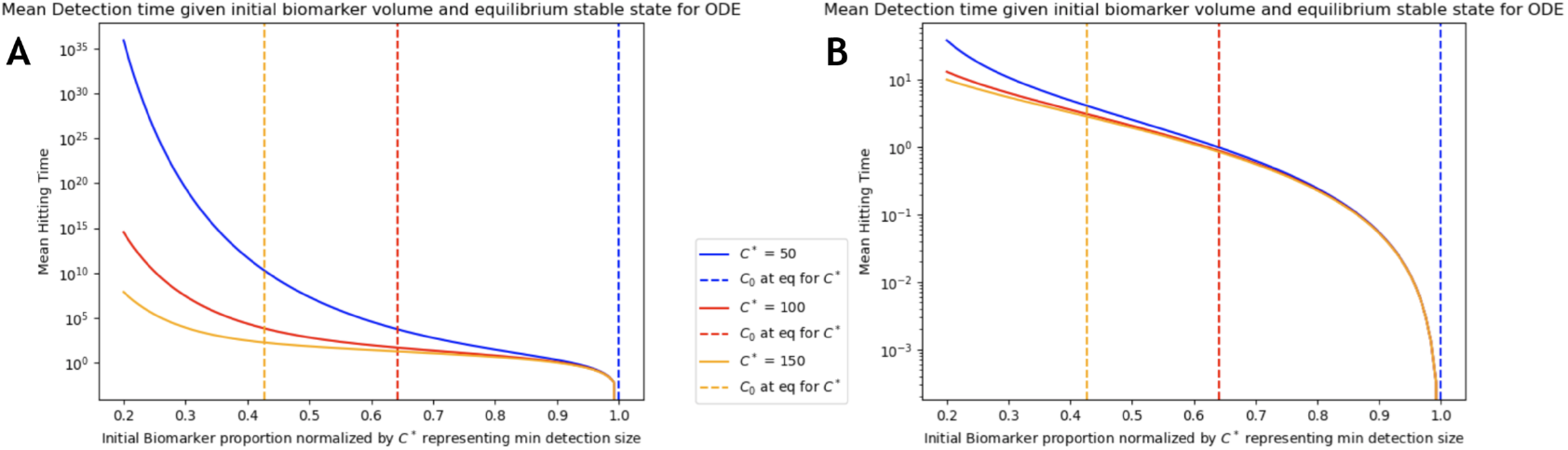
Log-scale Mean Detection time for heterogeneous immune mediated coexistence A) *σ*^2^ < 2*η*, B) *σ*^2^ > 2*η*. (*η* = 0.1, *σ* = 0.1, 0.45 respectively)

**Fig. 5.**
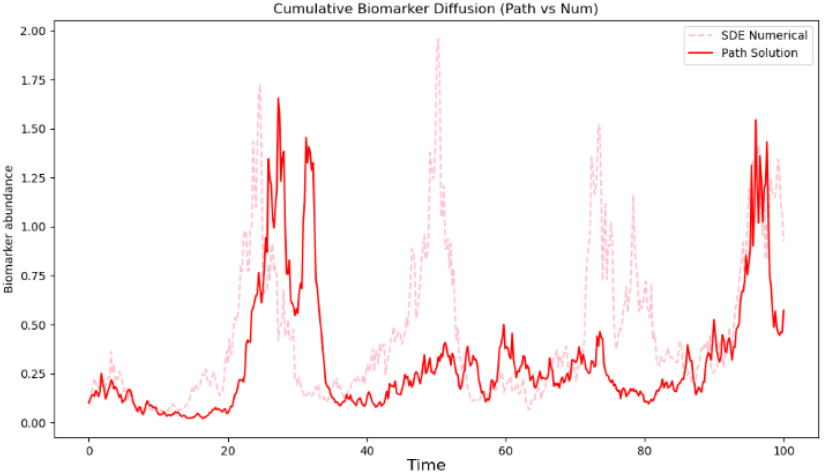
Biomarker stats with volatility parameter exceeding the biomarker degradation rate (*σ*^2^ > 2*η* with *σ* = 0.45, *η* = 0.1). This parametrization leads to earlier mean readout times.

These results suggest that therapeutic strategies which amplify volatility—either through modulation of immunedriven cell death variability or stochasticity in shedding mechanisms—may enhance detectability of dormant or early-stage tumors. Such approaches could improve early diagnosis by effectively accelerating escape from sub-threshold states.

Ideally, the mean time to detection serves as a statistic to see the impact of the underlying tumor-immune interaction to observed data. Given a well fitted underlying model, one could theoretically obtain the mean time to observe a readout given underlying dynamics, and compare to actual time to readout. This would provide one metric of the volatility describing the degree of biomarker source dependency given just these two release mechanisms for biomarker release.

### A.6. Anticipative approach to Biomarker Trajectory

Due to the lower detection limit of the biomarker, any observed signal indicates that the biomarker activity has crossed a critical threshold. In the previous section, we examined the stopping time at which the ratio *c* ^*^ reaches 1. In this section, we reverse the perspective: we assume that the observed signal, which occurs at some terminal time *T*, is instead interpreted as known at the initial time *t* = 0. Without loss of generality, we take *T* = 1. This forward-looking perspective, informed by future information, is analogous to techniques used in financial mathematics to model insider trading (45–47). We apply the same conceptual approach to the biomarker compartment. Specifically, we model this anticipative behavior using a generalized Itô integral. To that end, we employ anticipative stochastic calculus via the Ayed–Kuo stochastic integral (48). A brief introduction on the Ayed–Kuo integral is provided in S5.4 of the SI. We make a strong assumption that the observed signal received is a continuous functional of the noise at the terminal time, *ψ* (*W*1). This allows us to view the Eq. 2 as a linear stochastic differential equation with anticipating initial conditions.

*Theorem 7:* Given the following stochastic differential equation with anticipating initial condition,

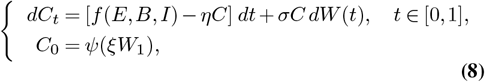

where *f* is defined as earlier in Table 4 and *ψ* 𝕔^2^(ℝ). Then, the solution in probability is given by

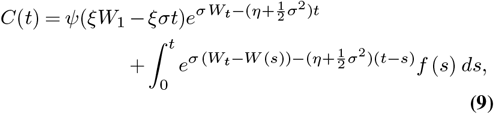

for *t* ∈ [0, 1].

*Proof:* Section S5.3 of the SI.

When we compare this result to the non-anticipating case in Eq. (4), the solution has a correction term scaled by the volatility, arising from the initial anticipating conditions and the driver Brownian motion. In the anticipating case, we initialize with the final information such as final biomarker levels. This retroactive modeling creates bias in the integral — and the correction term precisely removes that bias to maintain consistency with stochastic calculus rules. This means that although the anticipative formulation allows us to start from a future-informed initial state (such as *C*0 = *ψ* (*ξ W*1)), the resulting solution must still obey the structural integrity of stochastic integrals, particularly in preserving near-martingale properties, the general Itô formula and ensuring well-definedness under the anticipative integral. The correction term, therefore, acts as a compensator that restores the solution’s validity within the broader framework of stochastic analysis.

## B. Discussion

Challenges in tracking cancer dynamics near minimal detectable disease levels currently limit early diagnosis and identification of progression during disease modeling. To address this, we developed a general stochastic modeling framework to track the noisy biomarker release dynamics released from a population undergoing cell turnover and immune-mediated apoptosis. Our model provides a theoretical framework for understanding biomarker trajectories that depend on different release mechanics from the underlying tumor-immune interactions. We do so by coupling a system of ordinary differential equations representing a competitive heterogeneous tumor population interacting with the adaptive immune system with a family of stochastic differential equations representing the biomarker trajectories.

In this study, we used a one predator - two prey competitive Lokta-Volterra model where the prey (tumor cells) compete with each other under a shared carrying capacity. We show the underlying ODE model allows for coexistence between immune-targeted and immune-evasive cancer subpopulations. Relative altruism and egoism between these subpopulations play a critical role in this model, which suggests that the dynamics of tumor escape are shaped not just by individual subpopulation fitness, but also by their relative influence on one another. This finding highlights the importance of cooperation and competition as fundamental forces in shaping disease heterogeneity following immune escape. For example, cooperative behavior, such a shared secretion of growth factors, immune-modulating cytokines, or extracellular matrix remodeling enzymes, can enable certain subpopulations to persist despite being less fit in isolation (49). Conversely, competitive strategies, including resource monopolization or immune suppression, may allow one subpopulation to dominate but at the expense of overall tumor heterogeneity and adaptability (50, 51). In our model, a single-population escape, for instance, is only stable if the competing subpopulation behaves altruistically, acting as a cooperative partner by sharing resources or by absorbing immune pressure without reciprocation. In contrast, immune escape and stable coexistence of immune-evasive and immune-targeted populations requires that both populations adopt the same strategy, either mutually egoistic or mutually altruistic. If not, the egoistic subpopulation inevitably dominates, reducing heterogeneity to either an immune-evasive or targetable disease.

The complex interplay between model parameters and competitive dynamics is especially evident in the coexistence states. In the case of homogeneous immune-mediated coexistence, stability hinges on how these parameters align. Specifically, in Eq. (3), we derive a lower bound on the level of “assistance” required from the dying subpopulation for the dominant one to maintain balance with the immune system. This threshold depends on both the carrying capacity of the dominant subpopulation and the relative growth advantage between the two subpopulations. It effectively sets a minimum cooperation level needed to sustain equilibrium, highlighting how even a regressing subpopulation can play a stabilizing role in the tumor-immune landscape. This may imply that the extinction of a vulnerable subpopulation may reduce immune detection or competition for resources, indirectly supporting the survival of the more evasive subpopulation.

Recent work has investigated such behavior in breast cancer (30), wherein an altruistic subpopulation was observed to benefit the growth of the total population. These insights have potential implications for immunotherapy, as treatments that unintentionally eliminate the altruistic subpopulation may destabilize the system and pave the way for immune-resistant escape variants to expand unchecked. Additional experimental and modeling studies are needed to fully explore this therapeutic possibility.

Our model predicts that heterogeneous tumor escape requires additional constraints related to balancing the relative sub-population fitness values with the predication dynamics of the immune system. Our results predict a critical bound in the competitive parameter that partitions cooperative and competitive cancer strategies, which suggests that subtle shifts in competition for cancer populations operating close to this bound can dramatically reshape tumor-immune dynamics. Biologically, this result suggests that cooperative behaviors among tumor cells, including shared production of growth factors or immune modulators, may be under selective pressure not just for their immediate benefit, but for their role in maintaining population heterogeneity during immune surveillance. Moreover, detecting such cooperation could offer early clues about impending immune escape in the setting of immune control, and disrupting these interactions could be one targeting strategy to destabilize the tumor ecosystem. Our findings echo broader ecological principles in evolutionary theory, where population stability depends on the payoff structure of cooperative versus selfish strategies. In the context of cancer, this underscores that tumor progression is not merely a Darwinian race among isolated subpopulations, but also involves dynamic interactions between subpopulations exhibiting interdependent behavior (52).

The primary advantage of the foundational construction of the underlying model is that we obtain mathematical properties for the solutions of the system of equations. Using those features, we constructed a family of linear SDE’s to represent the biomarker trajectories from different release mechanisms. Here, we assume that biomarkers arise from inherent T cell death or from apoptosis arising from tumor-immune interaction. ct-DNA is one motivating example, but because we distinguish these sources, such a model could be used to more generally track independent signals arising from distinct cancer elimination modes. In the framework considered here, these stochastic trajectories depend on the tumor-immune interactions and competition between tumor cells under shared carrying capacity. This model allows us to obtain analytically tractable path-wise trajectories that exhibit both multiplicative behavior and log normal distribution and as such, the resulting behavior is more in line with the assumptions on biomarker release (36, 53). We also considered alternate models following the CIR process, but in this case, the nonlinear variant does not provide path-wise solutions crucial for the anticipative solution. In addition, non-negativity is not guaranteed while the variability for the CIR model is gamma-like rather than lognormal which might lead to underestimation of variance at high biomarker levels.

Our model represents heterogeneity through two subpopulations with different levels of immune targeting. While this abstraction captures the essential dynamics between more and less immune-targeted groups, real tumors span a broader distribution of immune recognition levels in addition to increasing clonal diversity. This simplification is compounded by the assumption that each SDE is driven by an independent Brownian motion, meaning fluctuations in one subpopulation’s biomarker shedding are unrelated to those in another. In practice, systemic influences could introduce correlated noise, potentially reshaping variance structure, modifying stability properties, and complicating parameter identifiability. In the anticipatory setting, we assume that the future information is a continuous functional of the Brownian motion. This requirement ensures the mathematical validity of anticipative stochastic calculus but may be restrictive biologically, as real measurements can be noisy, discontinuous, or influenced by factors outside the modeled stochastic drivers.

Tumor escape states, associated with immune elimination, are characterized by elevated necrotic biomarker levels, while immune-mediated coexistence yields a sustained but lower apoptotic signal due to ongoing tumor–immune interactions. However, only the escape dynamics consistently surpass critical detection thresholds, whereas the coexistence state often remains sub-threshold despite its biological activity. This underscores a key limitation of detection strategies that rely solely on biomarker magnitude. Volatility emerges as a critical factor in modulating detectability. As indicated by the variance expression in Eq. (6), high volatility relative to the biomarker degradation rate can lead to earlier threshold-crossing events. In regimes where mean dynamics fall below detection limits, stochastic fluctuations alone may drive observability as seen in Fig. 4.

These findings indicate that therapeutic strategies aimed at increasing volatility—whether by enhancing the variability of immune-mediated cell death or by introducing greater stochasticity in biomarker shedding—could improve the detectability of early-stage or dormant tumors. By amplifying fluctuations, such approaches may facilitate earlier diagnosis by promoting more rapid emergence from otherwise undetectable sub-threshold states.

Finally, we consider an anticipative stochastic integration framework to offer a complementary perspective on biomarker dynamics. By conditioning the system on future biomarker measurements, this approach formally incorporates future information. While such anticipative formulations depart from standard adapted Itô framework, they can offer bias-corrected trajectories that reflect the influence of future events on the interpretation of current dynamics. This perspective may be especially relevant for post hoc analysis of biomarker data, where retrospective fitting implicitly assumes access to future information.

Unlike traditional stopping time analyses, which halt simulation at a data-dependent threshold (e.g., a biomarker crossing a detection limit), the anticipative formulation allows us to reconstruct plausible entire trajectories given partial or endpoint information. This can be particularly valuable for retrospective modeling, imputation, or path-based inference where early or sparse data are supplemented by later observations. While the theory supports existence and structure of such solutions, developing a tractable simulation algorithm for anticipative integrals remains an open problem and promising direction for future work (47, 54). In parallel, comparing the insights derived from anticipative paths versus stopping-time-constrained simulations would clarify when each framework offers the most utility—whether for prediction, diagnostics, or intervention design.

## ACKNOWLEDGEMENTS

JTG was supported by the Cancer Prevention Research Institute of Texas (RR210080) and the National Institute of General Medical Sciences of the NIH (R35GM155458). JTG is a CPRIT Scholar in Cancer Research.

