## Supplementary Information for "Coupled SDE-ODE Modeling of Tumor-Immune Dynamics to Infer Biomarker Release"

August 15, 2025

#### S1 Overview

Effective cancer therapies, while continuously improving, are often limited by lower detection limits of disease burden. In the setting of the cancer immunotherapy, tumor immune dynamics in this limit present the most pressing knowledge gap as cancer ultimate escape or elimination are often determined following an intervening period of population equilibrium sustained at low population size. Population dynamics in this small-population limit are inherently affected by intrinsic noise in the tumor-immune interaction, as are estimates of population disease burden by extrinsic noise in acquiring such estimates through associated biomarkers. Since tumor progression often involves a heterogeneous population of cells having variable fitness, the population dynamics that incorporate both immune-recognized and immune-evasive subpopulations play an important role in prognosticating disease outcome. Here, we present a modeling framework that investigates the dynamic interactions between tumor cells and the immune system, focusing on how these interactions influence biomarker levels. The framework combines deterministic elements, which describe tumor growth and immune responses, with stochastic components that capture the inherent variability in biomarker release. We simulate a heterogeneous tumor-immune interaction by including two distinct cancer populations: one having baseline immunogenicity and the other with enhanced immune evasion. Here, competition between subpopulations in the setting of differential immune selective pressures drive diverse tumor growth dynamics.

We develop a foundational theoretical framework to analyze the equilibrium dynamics of all three compartments. The model incorporates competitive interactions between cancer subpopulations, with parameters reflecting altruistic and egoistic behaviors based on how the subpopulations impact each other's growth in the setting of a shared carrying capacity. Our findings suggest that these competitive parameters are crucial in determining the stability of "winners" and "losers" in tumor competition. Such modeling assumptions can be viewed as analogs to asymmetric competition in the literature [1]. We hypothesize that altruistic and egoistic behaviors among cancer cells may correspond to immune-recognized and immune-escape variants [2]. To capture the extrinsic variability of biomarkers in circulation (in addition to intrinsic stochasticity at low-population sizes), we use a stochastic differential equation (SDE) model, where biomarker release rates are influenced by the tumor-immune interactions. In this context, ct-DNA is used as a motivating example, but our framework can be adapted to study the release dynamics of other biomarker signatures resulting from tumor-immune interaction.

The rest of the work is structured as follows. Sec. S2 describes an overview of the background and motivation for this model. Sec. S3 frames the general mathematical problem of interest. Sec. S4 discusses the dynamical behavior of the competitive Lotka-Volterra two-prey one-predator model. Equilibrium and stability analysis is performed. Sec. S5 discusses the stochastic portion of the model and mean and variance trajectories is obtained for different biomarker mechanics for the competing tumors. Hitting time analysis for mean time to detection is performed.

In the special case where there is one shared tumor population targeted uniformly by the immune system, we find that the dynamics of the system reduce to the single-predator, single-prey system. Interestingly, another special case, where both tumor subpopulations are jointly neither altruistic nor egoistic, enables an interpretation of the two tumor populations as one homogeneous population with cumulative growth and death rates.

#### S2 Background and Motivation

The interactions between cancer cells and the immune system play an important role in determining the progression of cancer and the response to therapeutic interventions. These interactions are highly complex, with tumor cells exhibiting dynamic behaviors like immune evasion, ecological competition, and phenotypic adaptation. As such, understanding these processes is crucial for developing effective cancer treatments. While much of the current literature on tumor-immune interactions has focused on deterministic models of tumor growth, there is a growing need for approaches that also incorporate the stochastic nature of cancer progression. Such approaches are useful for describing the inherent randomness within cancer population dynamics in addition to accounting for uncertainty inherent in measuring relevant biomarkers.

Circulating tumor DNA (ct-DNA) is one particular example of a biomarker for which sparse events are sampled and defines a lower limit of detection during cancer modeling [3].Ct-DNA consists of fragmented DNA released into the bloodstream by dying tumor cells, primarily through apoptosis and necrosis [4]. Because ct-DNA levels reflect tumor burden and disease dynamics, they provide valuable insights into cancer progression and treatment efficacy, particularly when monitoring minimal residual disease following therapy. However, capturing ct-DNA dynamics requires models that can account for both the deterministic aspects of tumor growth and immune interaction, as well as the stochastic variations in biomarker release driven by biological noise. Previous stochastic modeling approaches have been introduced to address the variability in ct-DNA dynamics, often relying on simplified tumor growth frameworks such as birth-death processes or agent-based models [5–8]. While these models can effectively represent tumor dynamics, they tend to overlook the critical role of immune interactions in influencing ct-DNA shedding. These approaches do not capture the full complexity of tumor-immune interactions, particularly in the setting of intra-tumor heterogeneity, nor do they fully account for the variability in biomarker release due to immune-driven tumor cell death or other stochastic processes.

In this work, we introduce a generalizable framework that models the complex interactions between tumor biology and the immune system, integrating both deterministic and stochastic elements. Our approach uses a system of ordinary differential equations (ODEs) to represent the tumor-immune dynamics, where tumor growth is limited by a shared carrying capacity and immune cells act as predators on the tumor. Specifically, we model two tumor subpopulations, one exhibiting greater immune evasion than the other, and allow each subpopulation to interact dynamically with the immune system. The classic Lotka-Volterra description of predator-prey dynamics is adapted to capture the competitive interactions between each subpopulation and their interactions with immune cells [9–12].

The immune system's role in controlling tumor growth is reflected in the tumor-immune interaction terms in the ODEs. These interactions lead to tumor cell death through apoptosis. The resource competition between the subpopulations under the shared carrying capacity drives another cell death mechanism via necrosis. Both of these cell death processes contribute to the release of a tumor biomarker. We focus on applying our model to ct-DNA specifically, wherein we frame ct-DNA dynamics using a stochastic differential equation (SDE). This SDE framework accounts for the variability in ct-DNA release due to these dynamic tumor-immune interactions, as well as inherent biological noise, such as DNA degradation and clearance from circulation [4]. By coupling the ODE model for tumor-immune dynamics with the SDE for biomarker release, we can capture the fluctuations in biomarker levels driven by both stochastic tumor growth and exogenous sampling noise.

The framework introduced here is highly adaptable and can be applied to a variety of biomarker contexts, though we focus our application on ct-DNA as one particular example. The model provides insights into how immune dynamics influence detectable biomarker levels, offering a more realistic portrayal of tumor dynamics and the underlying inferred dynamics. In particular, this approach allows for the examination of tumor burden and the effectiveness of therapeutic interventions in real time, while also incorporating the stochastic variability inherent to biomarker measurements. By integrating tumor-immune interactions and biomarker dynamics, our model provides a more comprehensive description of cancer progression and monitoring, immune response, and treatment outcomes, and it can be extended to other biomarkers with similar underlying mechanisms.

#### S3 Model Development

We introduce a general framework for modeling the dynamical interactions between cancer cells and the immune system. This approach captures both deterministic elements — such as cancer growth and immune response — and stochastic elements, which describe the inherent variability in biomarker release influenced by these interactions. In particular, the model centers on two cancer subpopulations, one of which exhibits greater immune evasion than the other, interacting dynamically with the adaptive immune system. Cells in each subpopulation grow under a shared carrying capacity, and the immune system acts as a predator for both cell subtypes. We assume that the dynamics are dominated by cancer cell division and immune recognition and omit the possibility of phenotypic transition between each cancer subpopulation.

To represent the tumor-immune interaction, we use a system of ordinary differential equations (ODEs) that models tumor growth as logistic, limited by a shared carrying capacity $K$. This approach distinguishes the cancer subpopulations based on their immune susceptibility, where immune cells effectively act as predators on each tumor compartment. Let $E$, $B$, and $I$ represent the evasive cancer subpopulation, basal immune-recognizable cancer subpopulation, and immune cell compartments, respectively. The relationships are given by the following system of ODEs:

$$\frac{dE}{dt} = rE\left(1 - \frac{E + pB}{K}\right) - \alpha EI, \tag{S1}$$

$$\frac{dB}{dt} = \gamma B\left(1 - \frac{qE + B}{K}\right) - \beta BI, \tag{S2}$$

$$\frac{dI}{dt} = aEI + bBI - \delta I, \tag{S3}$$

with initial conditions $E(0), B(0), I(0) > 0$ and model parameters $r, \gamma, \alpha, \beta, a, b, \delta > 0$. Here:

- $K$: Carrying capacity for the total cancer population,
- $r, \gamma$: Growth rates of the evasive and basal recognizable tumors, respectively,
- $\alpha, \beta$: Death rates due to tumor-immune interactions ($\alpha < \beta$),
- $a, b$: Immune cell growth rates triggered by interactions with tumor cells,
- $\delta$: Immune cell death rate,
- $p, q$: Parameters that scale the competitive effects between the tumor populations.

The competitive scaling parameters, $p$ and $q$, characterize each tumor subpopulation as either *altruistic* or *egoistic* based on their interactions under shared carrying capacity. For example, if $0 < p < 1$, the evasive tumor reduces the baseline tumor's effective growth rate (egoistic), whereas $p > 1$ implies an altruistic effect, allowing the baseline compartment greater growth within the carrying capacity. To model biomarker levels in circulation, such as ct-DNA, we use a stochastic differential equation (SDE) to capture both deterministic influences from tumor-immune interactions and inherent biological variability. Namely, we model biomarker levels, denoted by $C$, with the SDE:

$$dC = [f(E, B, I) - \eta C]\, dt + \sigma C dW(t), \tag{S4}$$

where, for $C(0) > 0$, $\sigma$ and $\eta$ represent the volatility and decay rates of the biomarker, respectively, and $W(t)$ represents the standard Brownian motion starting at zero. Here, $f(E, B, I)$ denotes different release mechanisms. For foundational understanding, we focus on ct-DNA as the biomarker in question. ct-DNA is released through apoptosis and necrosis, with rates influenced by tumor burden and immune interactions [4]. We assume that apoptotic shedding is linked to tumor-immune interactions, while necrotic shedding scales with tumor size. These assumptions then inspire the following forms of deterministic drift in Eq. (S4):

- **Evasive Tumor Biomarker Trajectory**: $f(E,B,I) = rE\frac{E+pB}{K} + \alpha EI$,
- **Basal Recognizable Tumor Biomarker Trajectory**: $f(E,B,I) = \gamma B\frac{qE+B}{K} + \beta BI$,
- **Apoptotic Cells Biomarker Trajectory**: $f(E,B,I) = \alpha EI + \beta BI$,
- **Necrotic Cells Biomarker Trajectory**: $f(E,B,I) = \gamma B\frac{qE+B}{K} + rE\frac{E+pB}{K}$.

This structure captures variability in biomarker levels due to cell turnover, DNA degradation, and clearance from circulation, offering a realistic portrayal of biomarker dynamics influenced by tumor and immune cell interactions. By connecting biomarker dynamics to the tumor-immune interactions from the ODEs, this framework not only tracks tumor burden and treatment response but also accommodates stochastic fluctuations inherent to measuring biomarkers near the lower detection threshold limit. This framework, motivated by general principles of tumor-immune interactions, can extend beyond ct-DNA dynamics to other biomarkers impacted by similar biological interactions. Ct-DNA is presented here as a specific application, illustrating how the model's flexibility and stochastic component can adapt to other biomarker contexts, allowing for a broad exploration of biomarker release mechanisms that are modulated by tumor and immune system dynamics.

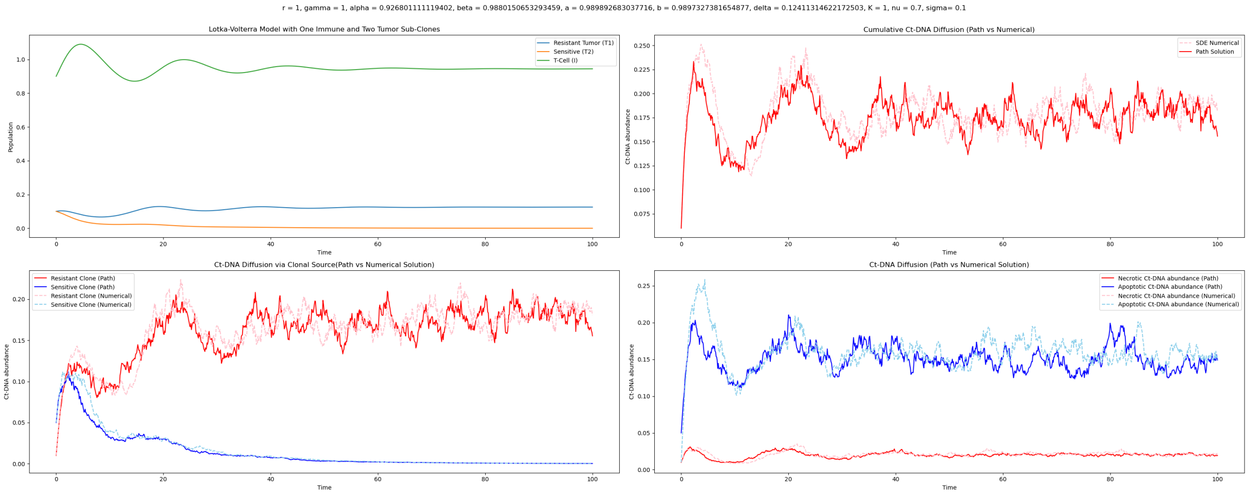


Figure S1: Lokta Volterra Pred-Prey Model for tumor-immune interaction. SDE simulated via both analytical path solution and numerical solution

### S4 Dynamical Behavior of the ODE

Let us consider the following system of equations in (S5). Namely,

$$\begin{aligned} dE &= rE(1 - \frac{E + pB}{K}) - \alpha EI \\ dB &= \gamma B(1 - \frac{qE + B}{K}) - \beta BI \\ dI &= aEI + bBI - \delta I. \end{aligned} \tag{S5}$$

with $p, q \in (0, 1)$ and $E(0), B(0), I(0) \geq 0$.

#### S4.1 Existence/ Uniqueness

The right hand side of the system of equations given by S5 are all continuous and Lipschitz on $\mathbf{C}[0, T]$. Via the Picard-Lindeoff theorem we have the existence and uniqueness of the solutions in a small neighborhood around the initial conditions. Namely, there exists $\epsilon > 0$ such that the solution both exists and is unique on the interval $[t_0 - \epsilon, t_0 + \epsilon]$. Since our system is Lipschitzian on $[0, T]$, this result can be extended to the whole interval. Finally, the positive initial conditions imply that the solutions are non-negative.

#### S4.2 Equilibrium States and Stability

Here we investigate the existence and stability of the system at the equilibrium states. We will focus in the interior equilibrium states for a major part of the analysis. The equilibrium states are the population values where the population levels stabilize:

$$dE = dB = dI = 0. \tag{S6}$$

This implies that,

$$\begin{aligned} & rE(1 - \frac{E + pB}{K}) - \alpha EI = 0 \\ \iff & \left( r(1 - \frac{E + pB}{K}) - \alpha I \right) E = 0 \\ \iff & E = 0 \text{ or } I = \frac{r}{\alpha}(1 - \frac{E + pB}{K}) \end{aligned} \tag{S7}$$

Similarly,

$$\begin{aligned} & \gamma B(1 - \frac{qE + B}{K}) - \beta BI = 0 \\ \iff & \left( \gamma(1 - \frac{qE + B}{K}) - \beta I \right) B = 0 \\ \iff & B = 0 \text{ or } I = \frac{\gamma}{\beta}(1 - \frac{qE + B}{K}) \end{aligned} \tag{S8}$$

Finally,

$$\begin{aligned} & aEI + bBI - \delta I = 0 \\ \iff & I = 0 \text{ or } aE + bB = \delta \end{aligned} \tag{S9}$$

##### S4.2.1 Boundary Equilibrium States

We can view boundary states at "ultimate winners or losers" in this predator-prey game between the adaptive immune system and the tumor subpopulations. Clearly, the case where all populations are absent, $(0, 0, 0)$, is the trivial equilibrium solution. If both the tumor compartments are zero, $E = B = 0$ then for the equilibrium conditions (S6) to hold, $I = 0$. As such, we next consider cases where one of the tumor subpopulations is not identically zero.

We first assume that the baseline tumor population at equilibrium is non zero. Hence, $E = 0$ and $I = \frac{\gamma}{\beta}(1 - \frac{qE+B}{K})$. We have two ways to satisfy (S6):

**Case 1:** $I = 0$

$$I = \frac{\gamma}{\beta}(1 - \frac{qE + B}{K}) = 0$$
$$\iff B = K$$

Thus, the equilibrium state is at $(0, K, 0)$ where the immune system and the evasive subpopulation are eliminated and the baseline recognizable subpopulation grows to carrying capacity.

**Case 2:** $aE + bB = \delta$

$$aE + bB = \delta$$
$$\implies B = \frac{\delta}{b}$$

Using this value of $B$, we find that

$$I = \frac{\gamma}{\beta}(1 - \frac{qE + B}{K}) = \frac{\gamma}{\beta}\left(1 - \frac{\delta}{bK}\right).$$

This gives us the equilibrium state

$$\left(0, \frac{\delta}{b}, \frac{\gamma}{\beta}\left(1 - \frac{\delta}{bK}\right)\right) \text{ with } K > \frac{\delta}{b}.$$

Note that, this equilibrium state is representative of the case where the baseline recognizable tumor out-competes the evasive cells and strikes a balance with the immune compartment. The equilibrium state for baseline recognizable cells reflects the balance between the immune death rate and the subsequent growth rate via immune interaction with baseline cells while the equilibrium state for the immune compartment reflects the balance struck between the baseline tumor growth rate and the subsequent death rate imposed on the baseline recognizable cells via immune interaction scaled logistically with respect to how close the equilibrium state is to the carrying capacity. We can use the same logic to find the other two boundary equilibrium states for when the evasive cells out-competes the baseline cells ($B = 0$ and $E$ nonzero).

Finally, we look at the case where $I = 0$ with $E$, $B$ nonzero. Here,

$$0 = \frac{r}{\alpha}(1 - \frac{E + pB}{K}) \implies E + pB = K,$$
$$0 = \frac{\gamma}{\beta}(1 - \frac{qE + B}{K}) \implies qE + B = K.$$

Solving the above system of equations, we obtain

$$E = \frac{K\,(1-p)}{1-pq}, \qquad B = \frac{K\,(1-q)}{1-pq}.$$

Under these constraints for $p, q$ , we find that $pq \neq 1$ and that the solutions for the tumor compartments are non-negative and bounded above by the carrying capacity. This therefore represents the case where the immune system is eliminated while the tumor populations strike coexistence. We note that for such an equilibrium condition to exist, the tumor subpopulations would need to be jointly egoistic ($0 < p, q < 1$) or jointly alturistic ($p, q > 1$).

So summarize, we have the following boundary equilibrium states:

- $(0, 0, 0)$,
- $(0, K, 0)$,
- $(0, \frac{\delta}{b}, \frac{\gamma}{\beta}(1 - \frac{\delta}{bK}))$,
- $(K, 0, 0)$,
- $(\frac{\delta}{a}, 0, \frac{r}{\alpha}(1 - \frac{\delta}{aK}))$,
- $\left(\frac{K(1-p)}{1-pq}, \frac{K(1-q)}{1-pq}.0\right)$

###### S4.2.2 Interior States

We pivot to the interior state where $K \geq E, B, I > 0$ and (S6) holds. While the previous boundary conditions represented outcomes where at-least one population is eliminated, the interior state represents sustained co-existence between all three compartments. As such, we have that

$$
\begin{aligned}
\frac{r}{\alpha}(1 - \frac{E + pB}{K}) &= \frac{\gamma}{\beta}(1 - \frac{qE + B}{K}), \\
\iff r\beta(K - E - pB) &= \gamma\alpha(K - qE - B), \\
\iff (\gamma\alpha q - r\beta)\, E + (\gamma\alpha - r\beta p)\, B &= K\,(\gamma\alpha - r\beta)\,, \\
\iff \psi E + \phi B &= \theta,
\end{aligned} \tag{S10}
$$

where

$$\psi \triangleq \gamma\alpha q - r\beta \quad \phi \triangleq \gamma\alpha - r\beta p \quad \theta \triangleq K(\gamma\alpha q - r\beta).$$

For such an interior solution to exist, $\phi$ and $\psi$ can not be zero simultaneously. Thus,

$$\gamma\alpha q \neq r\beta \text{ and } \gamma\alpha \neq r\beta p \iff p \neq \frac{r\beta}{\gamma\alpha} \text{ and } q \neq \frac{\gamma\alpha}{r\beta}. \tag{S11}$$

Interestingly, this implies a similar condition for existence as we obtained in the tumor coexistence case: $pq \neq 1$. Indeed, if $p = q = 1$, the equilibrium conditions would imply that either at equilibrium, $I = 0$, which contradicts the interior solution case, or there exists a constant $k_0$ such that $\gamma\alpha = k_0 r\beta$. In the latter case, we find that $dE = k_0 dS$ which would mean that the evasive population is nothing but a scaled version of the baseline recognizable one. This also means that, under this very restrictive condition on the competitive rates, the two heterogeneous tumor populations can be modeled via cumulative tumor population given by $dT_{cum} = (1 + k_0)dT_{cum}$.

We use Eq. (S10) alongside the fact that $aE + bB = \delta$ at the interior equilibrium state to obtain

$$E = \frac{\delta\phi - \theta b}{\phi a - \psi b}, \quad B = \frac{a\theta - \delta\psi}{a\phi - b\psi}. \tag{S12}$$

The equilibrium state for $I$ can be found via Eq. (S7) or Eq. (S8). From our discussion on Section S4.1, we have that the solution is positive and finite. As such, we can use the form of the equilibrium state to find conditions for existence of the solution.

- **Finite Solutions:** In addition to the criteria given by Eq. (S11), we have that

$$\begin{aligned} &\phi a - \psi b \neq 0 \\ \iff\ & p \neq \frac{\gamma\alpha}{r\beta} + \frac{b}{a}\left(1 - \frac{\gamma\alpha q}{r\beta}\right) \end{aligned}$$

- **Well-Defined Solutions:** From the construction of the model itself, we have that $0 < E, B \leq K$. First, we address non-negativity and then move to the boundededness condition. From Eq. (S12), $\delta\phi - \theta b$, $a\theta - \delta\psi$ , and $\phi a - \psi b$ all have the same sign. This provides an avenue by which we may analyze parametric regimes for the competition parameters $p$ and $q$, thereby studying the effects of relative "egoism" and "alturism" in each cancer subpopulations. Without loss of generality, we assume all of the expressions to be negative with the understanding that the same calculations would apply for the other case but with an inequality switch. We look into the first term,

$$\begin{aligned} 0 &> \delta\phi - \theta b = (\gamma\alpha - r\beta p)\delta - K(\gamma\alpha - r\beta)b \\ \iff p &> \frac{\gamma\alpha}{r\beta} - \frac{bK}{\delta}\frac{\gamma\alpha}{r\beta} + \frac{bK}{\delta} \\ \iff p &> 1 + \left(\frac{bK}{\delta} - 1\right)\left(1 - \frac{\gamma\alpha}{r\beta}\right) \triangleq p_0. \end{aligned} \tag{S13}$$

For the second term, we have

$$\begin{aligned} 0 &> a\theta - \delta\psi = (\gamma\alpha - r\beta)aK - \delta(\gamma\alpha q - r\beta), \\ \iff q &> \frac{r\beta}{\gamma\alpha} + \frac{aK}{\delta} - \frac{aK}{\delta}\frac{r\beta}{\gamma\alpha}, \\ \iff q &> 1 + \left(\frac{aK}{\delta} - 1\right)\left(1 - \frac{r\beta}{\gamma\alpha}\right) \triangleq q_0. \end{aligned} \tag{S14}$$

Finally, for the last term, we get,

$$\begin{aligned} 0 &> \phi a - \psi b = (\gamma\alpha - r\beta p)a - (\gamma\alpha q - r\beta)b \\ \iff \frac{a}{b}p + \frac{\gamma\alpha}{r\beta}q &> \frac{a}{b}\frac{\gamma\alpha}{r\beta} + 1 \end{aligned} \tag{S15}$$

Summarizing the above discussion for the negative case, Eq. (S14) and (S14) act as lower bounds for how helpful both cancer subpopulations need to be for the interior equilibrium state to exist. The model parameter specifications allow one subpopulation to be more egoistic and the other more altruistic. Indeed, if the product of the death rate of the baseline recognized tumor and the growth rate of the evasive tumor is more than the product of the death rate of the evasive tumor and the growth rate of the baseline tumor, $r\beta > \gamma\alpha$, then $p_0 > 1 > q_0$, then the relative net growth of the evasive tumor is greater than that of the baseline tumor. For this equilibrium point, the evasive tumor must necessarily be altruistic ( $p > p_0 > 1$), while the baseline recognized tumor is allowed a little bit of egoism ($q \in (q_0, 1) \cup [1, \infty)$). The reverse behavior can be obtained from the positive case as well. The third condition then puts a lower (or upper) bound for the joint contribution. We note here that the interior solution would not exist if one subpopulation is egoistic and one altruistic with respect to the $p_0$ and $q_0$ values. Now, we pivot to the immune compartment. From Eq. (S7) and Eq. (S8), we have that at equilibrium,

$$\frac{r}{\alpha K}(K - E - pB) = I = \frac{\gamma}{\beta K}(K - qE - B).$$

Thus, we can see that non-negativity for $I$ requires that $E + pB < K$ and $qE + B < K$. This condition alongside the fact that the carrying capacity condition imposes $E, B \leq K$, implies that there exists upper bound values for both $p, q$, say $p_{max}, q_{max}$, independent of the $p_0, q_0$ values. In summary, for the interior state to to exist, we would need $p < p_{max}, q < q_{max}$ and one of the following to hold,

– either $p > p_0$ and $q > q_0$, or

– $p < p_0$ and $q < q_0$.

##### S4.2.3 Stability of Equilibrium States

**Local Stability:**
The system of ODE's described by Eq. (S5) is nonlinear. One way to check stability is to linearize the model around the equilibrium states. To perform this linearization, we calculate the Jacobian matrix by taking the partial derivatives of the system's equations with respect to each variable. These partial derivatives form part of the Jacobian matrix, which represents the linearized system near an equilibrium state. Analyzing the eigenvalues of this Jacobian allows us to assess the local stability of the equilibrium. How close is the linearized model to the actual non-linear model? From the Hartman-Grobman Theorem, we have that near a hyperbolic equilibrium state (one where the Jacobian has no eigenvalues with zero real parts), the behavior of the nonlinear system is topologically equivalent to its linearized version [13]. Taking the partials of the system given by (S5), we get

$$\begin{aligned} f(E,B,I) &\triangleq rE(1-\frac{E+pB}{K})-\alpha EI \\ g(E,B,I) &\triangleq \gamma B(1-\frac{qE+B}{K})-\beta BI \\ h(E,B,I) &\triangleq aEI+bBI-\delta I. \end{aligned} \tag{S16}$$

For $f$,

$$\begin{aligned} f_E &= r\left(1-\frac{2E+pB}{K}\right)-\alpha I, \\ f_B &= -\frac{rpE}{K}, \\ f_I &= -\alpha E. \end{aligned}$$

For $g$,

$$\begin{aligned} g_E &= -\frac{\gamma qB}{K}, \\ g_B &= \gamma\left(1-\frac{qE+2B}{K}\right)-\beta I, \\ g_I &= -\beta B. \end{aligned}$$

For $h$,

$$\begin{aligned} h_E &= aI, \\ h_B &= bI, \\ h_I &= aE+bB-\delta. \end{aligned}$$

Then the Jacobian of the system is given by

$$J=\begin{bmatrix} r\left(1-\frac{2E+pB}{K}\right)-\alpha I & -\frac{rpE}{K} & -\alpha E \\ -\frac{\gamma qB}{K} & \gamma\left(1-\frac{qE+2B}{K}\right)-\beta I & -\beta B \\ aI & bI & aE+bB-\delta \end{bmatrix}$$

**Boundary Equilibrium States:**

- $e_0 = (0, 0, 0)$ :

$$J\Big|_{e_0} = \begin{bmatrix} r & 0 & 0 \\ 0 & \gamma & 0 \\ 0 & 0 & -\delta \end{bmatrix}$$

  The Jacobian at $e_0$ has eigenvalues $r > 0$, $\gamma > 0$, and $-\delta < 0$, indicating that this state is unstable. The positive eigenvalues for the evasive and baseline tumor subpopulations imply that near zero population, both tumor subpopulations will grow rather than die off. The negative eigenvalue shows that the immune system is suppressed in this region. This instability suggests that the immune system is unable to control tumor growth, and small perturbations will lead to the expansion of tumor populations, meaning joint tumor extinction is unlikely.

- **Baseline tumor wins:** $e_1 = (0, K, 0)$ :

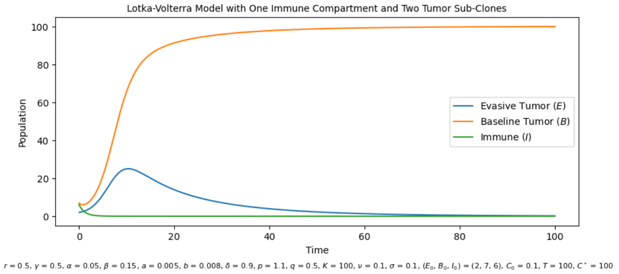


Figure S2: Simulated Baseline Win Scenario with params given. Parameter choices provided at the bottom of the figure

$$J\Big|_{e_1} = \begin{bmatrix} r(1-p) & 0 & 0 \\ -\gamma q & -\gamma & -\beta K \\ 0 & 0 & bK - \delta \end{bmatrix}$$

  The Jacobian at $e_1$ has eigenvalues $\lambda = -\gamma, r(1-p), bK - \delta$. Here, $-\gamma < 0$, meaning that the local stability of the case where the baseline tumor wins depends the signs of the other two eigenvalues.Thus, $e_1$ is stable when the equilibrium value of the baseline tumor population is more than the carrying capacity and the evasive tumor population is altruistic.

  **Stability Condition:** $K < \frac{\delta}{b}$ and $p > 1$.

- **Evasive tumor wins:** $e_2 = (K, 0, 0)$ :

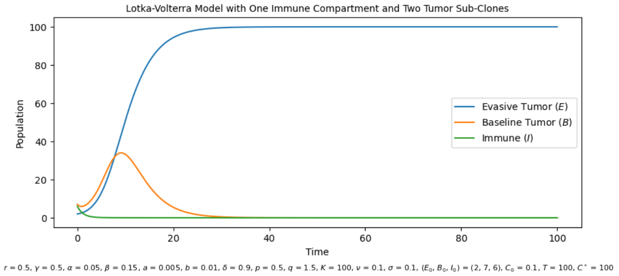


Figure S3: Simulated evasive Win scenario with given params. Parameter choices provided at the bottom of the figure

$$
J\Big|_{e_2} = \begin{bmatrix} -r & -rp & -\alpha K \\ 0 & \gamma\,(1-q) & 0 \\ 0 & 0 & aK-\delta \end{bmatrix}
$$

This is similar to the baseline winning case. The Jacobian at $e_1$ has eigenvalues $\lambda = -r,\ \gamma\,(1-q)\,,\ aK-\delta$. Thus, $e_2$ is stable when the equilibrium value of the evasive tumor population is less than the carrying capacity and the baseline tumor population is altruistic.

**Stability Condition:** $K < \frac{\delta}{a}$ and $q > 1$.

- **Evasive tumor loses, baseline tumor coexists with immune:** $e_3 = \left(0, \frac{\delta}{b}, \frac{\gamma}{\beta}\left(1 - \frac{\delta}{bK}\right)\right)$ :

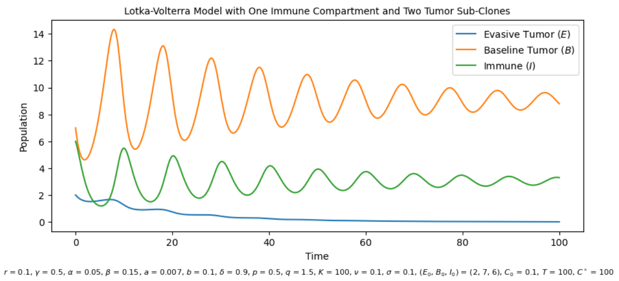


Figure S4: Simulated Evasive Loser Scenario with params given. Parameter choices provided at the bottom of the figure

$$
J\Big|_{e_3} = \begin{bmatrix} r\left(1 - \frac{p\delta}{bK}\right) - \frac{\gamma\alpha}{\beta}(1 - \frac{\delta}{bK}) & 0 & 0 \\ -\frac{\gamma q\delta}{bK} & -\frac{\gamma\delta}{bK} & -\beta\frac{\delta}{b} \\ \frac{a\gamma}{\beta}(1 - \frac{\delta}{bK}) & \frac{b\gamma}{\beta}(1 - \frac{\delta}{bK}) & 0 \end{bmatrix}
$$

The Jacobian at $e_3$ has eigenvalues

$$
\lambda = r\left(1 - \frac{p\delta}{bK}\right) - \frac{\gamma\alpha}{\beta}(1 - \frac{\delta}{bK})
$$

and $\lambda_\pm$, which satisfies the quadratic equation,

$$
\lambda^2 + \frac{\gamma\delta}{bK}\lambda + \delta\gamma(1 - \frac{\delta}{bK}) = 0.
$$

For the quadratic equation to have negative real parts, we use the Routh-Hurwitz Stability criterion [14] for quadratic polynomials which states that a monic quadratic equation will have negative real roots iff the coefficients are positive. This hold true as the baseline tumor equilibrium value is less than the carrying capacity for this equilibrium state. Finally, we need

$$
\begin{aligned}
r\left(1 - \frac{p\delta}{bK}\right) &< \frac{\gamma\alpha}{\beta}(1 - \frac{\delta}{bK}) \\
\iff rbK\beta - r\delta\beta p &< \gamma\alpha bK - \gamma\alpha\delta \\
\iff \delta(\gamma\alpha - r\beta p) &< bK(\gamma\alpha - r\beta)
\end{aligned}
$$

This is nothing but $\delta\phi - \theta b < 0$. Thus, from our interior equilibrium calculation in Eq. (S13), we find that the equilibrium state where the evasive tumor dies out while the baseline and the immune compartment coexist imposes a lower bound to how altruistic the the evasive population can be. Namely, we get the following stability criterion.

**Stability Criterion:**

- $\frac{\delta}{b} < K$
- $p > 1 + \left(\frac{bK}{\delta} - 1\right)\left(1 - \frac{\gamma\alpha}{r\beta}\right)$

- **Baseline tumor loses, evasive tumor coexists with immune:** $e_4 = (\frac{\delta}{a}, 0, \frac{r}{\alpha}(1 - \frac{\delta}{aK}))$ :

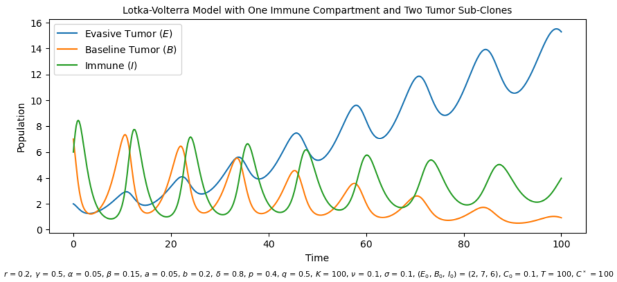


Figure S5: Simulated baseline Loser Scenario with params given. Parameter choices provided at the bottom of the figure

$$J\Big|_{e_4} = \begin{bmatrix} -\frac{r\delta}{aK} & -\frac{rp\delta}{aK} & -\alpha\frac{\delta}{a} \\ 0 & \gamma\left(1 - \frac{q\delta}{aK}\right) - \frac{r\beta}{\alpha}(1 - \frac{\delta}{aK}) & 0 \\ a\frac{r}{\alpha}(1 - \frac{\delta}{aK}) & \frac{rb}{\alpha}(1 - \frac{\delta}{aK}) & 0 \end{bmatrix}$$

With a similar argument to that of the previous case, we can say that the stability at this equilibrium value depends on the sign of the eiganvalue given by

$$\lambda = \gamma\left(1 - \frac{q\delta}{aK}\right) - \frac{r\beta}{\alpha}\left(1 - \frac{\delta}{aK}\right),$$

and the roots of the following quadratic equation

$$\lambda^2 + \frac{r\delta}{aK}\lambda + \delta r(1 - \frac{\delta}{aK}) = 0.$$

Similar to the symmetric case where the baseline coexists with immune compartment, we have that

$$\gamma\left(1 - \frac{q\delta}{aK}\right) - \frac{r\beta}{\alpha}\left(1 - \frac{\delta}{aK}\right) < 0 \iff a\theta - \delta\psi < 0$$

Using the result of Eq. (S14), we obtain a lower bound for how altruistic the baseline tumor needs to be in order for the evasive tumor to eliminate it and coexist with the immune compartment.

**Stability Criterion:**

- $\frac{\delta}{a} < K$
- $q > 1 + \left(\frac{aK}{\delta} - 1\right)\left(1 - \frac{r\beta}{\gamma\alpha}\right)$

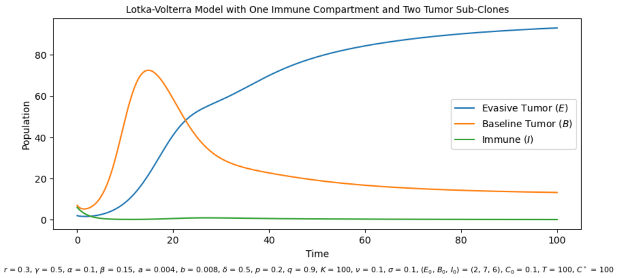


Figure S6: Simulated Evasive-immune co-existence with given params. Parameter choices provided at the bottom of the figure

- **Immune loses, tumor co-existence**: $e_5 = \left(\frac{K(1-p)}{1-pq}, \frac{K(1-q)}{1-pq}, 0\right)$

$$J\Big|_{e_5} = \begin{bmatrix} \frac{-rE}{K} & -\frac{rpE}{K} & -\alpha E \\ -\frac{\gamma qB}{K} & \frac{-\gamma B}{K} & -\beta B \\ 0 & 0 & aE + bB - \delta \end{bmatrix}$$

The Jacobian at $e_5$ has eigenvalues given by

$$\lambda = \delta - aE - bB$$

and the roots of the following quadratic equation

$$\lambda^2 + \frac{r(1-p) + \gamma(1-q)}{1-pq}\lambda + r\gamma\frac{(1-p)(1-q)}{1-pq} = 0.$$

From earlier discussion in Sec. S4.2.1, we have that the tumor co-existence equilibrium requires both tumor subpopulations to simultaneously be either both egoistic or both altruistic.

**Stability Criterion:**

- $p, q < 1$ or $p, q > 1$
- $\delta > \frac{K}{1-pq}\left(a(1-p) + b(1-q)\right)$

**Interior Equilibrium State:**

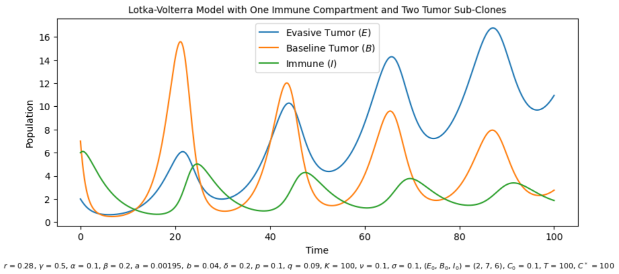


Figure S7: Simulated Co-existence dynamics with parameters given above

Now we look into the stability of the co-existence equilibrium state. While the previous boundry cases represented the “winners” and “losers” in this ecological game, the interior state represents where evasive, baseline and the imumune compartments exist while maintaining a balance with each other. Note that the Jacobian for this state is given by:

$$J\Big|_{e_6} = \begin{bmatrix} -\frac{r}{K}\frac{\delta\phi-\theta b}{\phi a-\psi b} - \lambda & -\frac{rp}{K}\frac{\delta\phi-\theta b}{\phi a-\psi b} & -\alpha\frac{\delta\phi-\theta b}{\phi a-\psi b} \\ -\frac{\gamma q}{K}\frac{a\theta-\delta\psi}{a\phi-b\psi} & -\frac{\gamma}{K}\left(\frac{a\theta-\delta\psi}{a\phi-b\psi}\right) - \lambda & -\beta\frac{a\theta-\delta\psi}{a\phi-b\psi} \\ a\frac{r}{\alpha K}\left(K - \frac{\delta\phi-\theta b}{\phi a-\psi b} - p\frac{a\theta-\delta\psi}{a\phi-b\psi}\right) & b\frac{\gamma}{\beta K}\left(K - q\frac{\delta\phi-\theta b}{\phi a-\psi b} - \frac{a\theta-\delta\psi}{a\phi-b\psi}\right) & a\frac{\delta\phi-\theta b}{\phi a-\psi b} + b\frac{a\theta-\delta\psi}{a\phi-b\psi} - \delta - \lambda \end{bmatrix}$$

with the characteristic functions given by

$$\lambda^3 + A\lambda^2 + B\lambda + C$$

where,

$$\begin{aligned} A =& \frac{r\hat{E}}{K} + \frac{\gamma\hat{B}}{K} - a\hat{E} - b\hat{B} + \delta \\ B =& \frac{r\gamma\hat{E}\hat{B}}{K^2} - \frac{ra\hat{E}^2}{K} - \frac{rb\hat{E}\hat{B}}{K} + \frac{r\delta\hat{E}}{K} - \frac{\gamma a\hat{E}\hat{B}}{K} - \frac{\gamma b\hat{B}^2}{K} + \frac{\gamma\hat{B}\delta}{K} + b\gamma\hat{B} - \frac{b\gamma q\hat{E}\hat{B}}{K} - \frac{b\gamma\hat{B}^2}{K} \\ & - \frac{rp\gamma q\hat{E}\hat{B}}{K^2} + ar\hat{E} - \frac{ar\hat{T}_1^2}{K} - \frac{arp\hat{E}\hat{B}}{K^2} \\ C =& \left(-\frac{r\gamma a\hat{E}^2\hat{B}}{K^2} - \frac{r\gamma b\hat{E}\hat{B}^2}{K^2} + \frac{r\gamma\delta\hat{E}\hat{B}}{K^2} + \frac{rb\gamma\hat{E}\hat{B}}{K} - \frac{rb\gamma q\hat{E}^2\hat{B}}{K^2} - \frac{rb\gamma\hat{E}\hat{B}^2}{K^2}\right) \\ & + \left(\frac{rp\gamma qa\hat{E}^2\hat{B}}{K^2} + \frac{rp\gamma qb\hat{E}\hat{B}^2}{K^2} - \frac{rp\gamma q\delta\hat{E}\hat{B}}{K^2} - \frac{a\beta r^2p\hat{E}\hat{B}}{\alpha K} + \frac{a\beta r^2p\hat{E}^2\hat{B}}{\alpha K^2} + \frac{a\beta r^2p^2\hat{E}\hat{B}^2}{\alpha K^2}\right) \\ & - \alpha\hat{E}\left(\frac{b\gamma^2q\hat{B}}{\beta K} - \frac{b\gamma^2q^2\hat{E}\hat{B}}{\beta K^2} - \frac{b\gamma^2q\hat{B}^2}{\beta K^2} - \frac{a\gamma r\hat{B}}{\alpha K} + \frac{a\gamma r\hat{E}\hat{B}}{\alpha K^2} + \frac{a\gamma rp\hat{B}^2}{\alpha K^2}\right) \end{aligned}$$

and

$$\begin{aligned} \hat{E} =& \frac{\delta\phi - \theta b}{\phi a - \psi b} \\ \hat{B} =& \frac{a\theta - \delta\psi}{a\phi - b\psi} \end{aligned}$$

Hence, from the Routh-Hurwitz stability criterion [14], we have that the above expression will have negative real part in their eigenvalues if and only if the following inequalities hold for the above coefficients:

- $A, C > 0$
- $AB - C > 0$

**Global Stability:**

In the previous section, we described stability in a local sense. Namely, that the stability analysis via the linearized system only describes the behavior of trajectories close to the equilibrium, not their behavior far from it. In contrast, global stability concerns the stability of the system across the entire state space, not just near specific equilibria. A globally stable equilibrium means that, regardless of the initial conditions, the system will eventually settle into the equilibrium state over time.

Now we look into global stability for the interior equilibrium state. We employ Lyapunov's second method of stability to find conditions for global stability. Consider the following functional given by

$$L(E, B, I) = P(E - \hat{E} - \hat{E}\ln\frac{E}{\hat{E}}) + Q(B - \hat{B} - \hat{B}\ln\frac{B}{\hat{B}}) + R(I - \hat{I} - \hat{I}\ln\frac{I}{\hat{I}}) \tag{S17}$$

where $P, Q, R$ are positive constants to be chosen later. This form of functional allows us to measure deviations from the equilibrium while allowing for easier tractability with the time derivative operator. These properties help in verifying the $L$ is a Lyapunov function. Indeed, $L(\hat{E}, \hat{B}, \hat{I}) = 0$ and using the fact that, $\ln x \geq x - 1$ for $x > 0$,

$$\begin{aligned} P(E - \hat{E} - \hat{E} \ln \frac{E}{\hat{E}}) &\geq P(E - \hat{E} + \hat{E}(1 - \frac{E}{\hat{E}}) \\ &= P(E - \hat{E} + \hat{E} - \hat{E}\frac{E}{\hat{E}}) \\ &= 0 \end{aligned}$$

Hence, $L(E, B, I) \geq 0.$. Now we differentiate $L$ with respect to time to get

$$\begin{aligned} \frac{dL}{dt} &= P\left(\frac{E - \hat{E}}{E}\right)\frac{dE}{dt} + Q\left(\frac{B - \hat{B}}{B}\right)\frac{dB}{dt} + R\left(\frac{I - \hat{I}}{I}\right)\frac{dI}{dt} \\ &\triangleq \mathrm{I} + \mathrm{II} + \mathrm{III} \end{aligned}$$

For ease of calculation, lets focus on each part separately.

$$\begin{aligned} \mathrm{I} &= P\left(\frac{E - \hat{E}}{E}\right)\frac{dE}{dt} \\ &= P\left(\frac{E - \hat{E}}{E}\right)\left(rE(\left(1 - \frac{E + pB}{K}\right) - \alpha EI\right) \\ &= P\left(E - \hat{E}\right)\left(r\left(1 - \frac{E + pB}{K}\right) - \alpha I\right) \\ &= P\left(E - \hat{E}\right)\left(r\left(1 - \frac{E + pB}{K}\right) - \alpha I - r\left(1 - \frac{\hat{E} + p\hat{B}}{K}\right) + \alpha \hat{I}\right) \\ &= P\left(E - \hat{E}\right)\left(-\frac{r}{K}\left(E - \hat{E}\right) - \frac{rp}{K}\left(B - \hat{B}\right) - \alpha\left(I - \hat{I}\right)\right) \\ &= -\frac{rP}{K}\left(E - \hat{E}\right)^2 - \frac{prP}{K}\left(E - \hat{E}\right)\left(B - \hat{B}\right) - \alpha P\left(I - \hat{I}\right)\left(E - \hat{E}\right) \end{aligned}$$

$$\begin{aligned} \mathrm{II} &= Q\frac{B - \hat{B}}{B}\frac{dB}{dt} \\ &= Q\left(B - \hat{B}\right)\left(\gamma\left(1 - \frac{qE + B}{K}\right) - \beta I\right) \\ &= Q\left(B - \hat{B}\right)\left(-\frac{\gamma q}{K}\left(E - \hat{E}\right) - \frac{\gamma}{K}\left(B - \hat{B}\right) - \beta\left(I - \hat{I}\right)\right) \\ &= -\frac{\gamma q Q}{K}\left(B - \hat{B}\right)\left(E - \hat{E}\right) - \frac{\gamma Q}{K}\left(B - \hat{B}\right)^2 - \beta Q\left(I - \hat{I}\right)\left(B - \hat{B}\right) \end{aligned}$$

$$\begin{aligned} \mathrm{III} &= R\frac{I - \hat{I}}{I}\frac{dI}{dt} \\ &= R\left(I - \hat{I}\right)(aE + bB - \delta) \\ &= aR\left(I - \hat{I}\right)\left(E - \hat{E}\right) + bR\left(I - \hat{I}\right)\left(B - \hat{B}\right) \end{aligned}$$

Thus,

$$\mathrm{I}+\mathrm{II}+\mathrm{III} = -\frac{rP}{K}\left(E-\hat{E}\right)^2 - \frac{\gamma Q}{K}\left(B-\hat{B}\right)^2 - \left(\frac{Prp}{K}+\frac{\gamma q Q}{K}\right)\left(E-\hat{E}\right)\left(B-\hat{B}\right)$$
$$+ (aR-\alpha P)\left(I-\hat{I}\right)\left(E-\hat{E}\right) + (bR-\beta Q)\left(I-\hat{I}\right)\left(B-\hat{B}\right)$$

If we select $P=p$, $R=\frac{\alpha p}{a}$, and $Q=\frac{b\alpha p}{a\beta}$ then the above expression can be simplified to

$$\mathrm{I}+\mathrm{II}+\mathrm{III} = -\frac{rp}{K}\left(E-\hat{E}\right)^2 - \frac{\gamma b\alpha p}{a\beta K}\left(B-\hat{B}\right)^2 - \left(\frac{rp^2}{K}+\frac{\gamma q b\alpha p}{a\beta K}\right)\left(E-\hat{E}\right)\left(B-\hat{B}\right)$$

This means that, $\frac{dL}{dt} < 0$ when either of the following conditions hold:

- $E > \hat{E}$ and $B > \hat{B}$ or,
- $E < \hat{E}$ and $B < \hat{B}$.

Thus, if the above conditions hold, $L$ is a Lyapunov function and the interior state is asymptotically stable in Lyapunov's sense. This means that trajectories that have the evasive and baseline tumor population jointly above or below the equilibrium state will stably converge to the interior equilibrium state asymptotically.

#### S4.3 Nondimensionalization of the Model

For ease of fitting to data, we also consider the nondimentionalized version of the model. Nondimensionalization of an ODE model helps reduce the number of parameters by combining them into dimensionless groups, thus simplifying the system. By eliminating units and resealing variables, the model becomes easier to analyze, as fewer distinct parameters need to be considered. This reduction is particularly useful for our case. We first scale the tumor populations by their carrying capacity. Furthermore, we assume that there is a reference $I_0$ that we can obtain empirically or as a parameter from the agent based model. Then,

$$\bar{E} = \frac{E}{K} \qquad \bar{B} = \frac{B}{K} \qquad \bar{I} = \frac{I}{I_0}. \tag{S18}$$

Now we scale time by the growth rate of the baseline recognizable tumor subpopulation. Namely, let $\tau = \gamma t$. Under these scaling, we have the scaled evasive tumor dynamics,

$$\begin{aligned}
\frac{d\bar{E}}{d\tau} &= \frac{1}{\gamma K}\frac{dE}{dt}, \\
&= \frac{1}{\gamma K}\left(rE\left(1-\frac{E}{K}-p\frac{B}{K}\right) - \alpha E I_0 \frac{I}{I_0}\right), \\
&= \frac{r}{\gamma}\bar{E}\left(1-\bar{E}-p\bar{B}\right) - \frac{\alpha I_0}{\gamma}\bar{E}\bar{I}.
\end{aligned}$$

For the baseline recognizable tumor subpopulation,

$$\begin{aligned}
\frac{d\bar{B}}{d\tau} &= \frac{1}{\gamma K}\frac{dB}{dt}, \\
&= \frac{1}{\gamma K}\left(\gamma B\left(1-q\frac{E}{K}-\frac{B}{K}\right) - \beta B I_0 \frac{I}{I_0}\right), \\
&= \bar{B}(1-q\bar{E}-\bar{B}) - \frac{\beta I_0}{\gamma}\bar{B}\bar{I}.
\end{aligned}$$

Finally, the immune compartment becomes

$$\begin{aligned}\frac{d\bar{B}}{d\tau} &= \frac{1}{\gamma I_0}\left(aEI + bBI - \delta I\right), \\ &= \frac{aK}{\gamma}\frac{E}{K}\frac{I}{I_0} + \frac{bK}{\gamma}\frac{B}{K}\frac{I}{I_0} - \delta\frac{I}{I_0}, \\ &= \frac{aK}{\gamma}\bar{E}\bar{I} + \frac{bK}{\gamma}\bar{B}\bar{I} - \frac{\delta}{\gamma}\bar{I}.\end{aligned}$$

Thus, we have a new system of equations that reflect the dynamics of our original model albiet with less parameters. Namely,

$$\begin{aligned}d\bar{E} &= \bar{r}\bar{E}\left(1 - \bar{E} - p\bar{B}\right) - \bar{\alpha}\bar{E}\bar{I}, \\ d\bar{B} &= \bar{B}(1 - q\bar{E} - \bar{B}) - \bar{\beta}\bar{B}\bar{I}, \\ d\bar{I} &= \bar{a}\bar{E}\bar{I} + \bar{b}\bar{B}\bar{I} - \bar{\delta}\bar{I},\end{aligned} \tag{S19}$$

where the re-scaled parameters are

$$\bar{r} = \frac{r}{\gamma}, \qquad \bar{\alpha} = \frac{\alpha I_0}{\gamma}, \qquad \bar{\beta} = \frac{\beta I_0}{\gamma}, \qquad \bar{a} = \frac{aK}{\gamma}, \qquad \bar{b} = \frac{bK}{\gamma}, \qquad \bar{\delta} = \frac{\delta}{\gamma}.$$

As a sanity check, we want to see if the parameters are unitless. Here,

$$\begin{aligned}&[r] = \text{time}^{-1}, \qquad [\gamma] = \text{time}^{-1}, \qquad [\alpha] = \text{population}^{-1} \times \text{time}^{-1}, \\ &[\beta] = \text{population}^{-1} \times \text{time}^{-1}, \qquad [a] = \text{population}^{-1} \times \text{time}^{-1} \qquad [b] = \text{population}^{-1} \times \text{time}^{-1} \\ &[\delta] = \text{time}^{-1}, \qquad [K] = \text{population}, \qquad [I_0] = \text{population}.\end{aligned}$$

Thus, we can see that the parameters and the system itself are both unit-less.

### S5 Dynamical Behavior of the SDE

Recall that we consider a system of SDE to model biomarker levels. Namely, $C(t), t \in [0, T]$ represents the biomarker levels in blood, normalized by the minimal size required for the tumor to be detected in liquid biopsies, and is given by the following class of SDE

$$dC = [f(E, B, I) - \eta C]\, dt + \sigma C\, dW(t), \qquad t \in [0, T]$$

where, $W(t)$ is the standard Brownian motion starting at zero. Here, $\sigma$ represents the random fluctuations in biomarker levels, which can be influenced by various factors such as dynamic changes in tumor shedding rates, blood circulation, and the detection process itself. On the other hand, $\eta$ represents the rate at which biomarker is cleared from the bloodstream. This can occur via natural processes like degradation by enzymes, clearance through renal pathways, or uptake by immune cells. Let $c^*$ be the minimal size required for the tumor. For foundational understanding, let us motivate the functional forms of the drift term by interpreting ct-DNA to be the biomarker to model. Under that assumption, we consider the possible forms and rationale for the drift term $f$:

- Evasive Tumor Biomarker Trajectory: $\frac{1}{c^*}\left(rE(\frac{E+pB}{K}) + \alpha EI\right)$,
- Baseline Tumor Biomarker Trajectory: $\frac{1}{c^*}\left(\gamma B(\frac{qE+B}{K}) + \beta BI\right)$,
- Apoptotic Cells Biomarker Trajectory: $\frac{1}{c^*}\left(\alpha EI + \beta BI\right)$,
- Necrotic Cells Biomarker Trajectory: $\frac{1}{c^*}\left(\gamma B(\frac{qE+B}{K}) + rE(\frac{E+pB}{K})\right)$.

Table S1 displays the parameter and compartment dependencies for each of the release mechanisms for evasive and baseline tumors.

| | **Apoptotic** | **Necrotic** |
|---|---|---|
| **Evasive** | $\frac{1}{c^*}(\alpha EI)$ | $\frac{1}{c^*}\left(rE(\frac{E+pB}{K})\right)$ |
| **Baseline** | $\frac{1}{c^*}(\beta BI)$ | $\frac{1}{c^*}\left(\gamma B(\frac{qE+B}{K})\right)$ |

Table S1: Form of $f$ and their rationale for Evasive and Baseline Tumors

For this text, we focus on the general class of linear stochastic differential equations with $f$ as listed above. We also consider the linear combination of the entries in Table S1. We note that, in simulations, we consider the same Brownian motion for evasive apoptotic and necrotic compartments while considering another independent Brownian motion for the two compartments in the baseline case.

#### S5.1 Properties of the Solution:

##### S5.1.1 Existence and Uniqueness of the Solution

We have that that $E$ and $B$ are both bounded via the logistic growth term. This implies that $\exists\, I \in [0, \infty)$ such that all the derivatives given by S5 are all negative. As such, $I$ is also upper bounded in the interval $[0, T]$. Since all the derivatives are bounded, we have that the functions, $E$, $B$, and $I$ are Lipschitz in time. As sums of products of Lipschitz functions, $f$ is Lipschitz in time. Thus, the unique solution of the system of SDEs given by S4 with $E(0), B(0), I(0) > 0$ and $C(0) > 0$ both exists (Theorem 11.1.1 of [15]) and given by

$$C(t) = C(0)e^{\sigma W(t)-(\eta+\frac{1}{2}\sigma^2)t} + \int_0^t e^{\sigma(W(t)-W(s))-(\eta+\frac{1}{2}\sigma^2)(t-s)} f\left(E(s), B(s), I(s)\right) ds \quad \text{(S20)}$$

Furthermore, by the construction of $f$ as sums and products of non-negative functions, $f$ itself is non-negative. This implies that $C(t) \geq 0$.

###### S5.1.2 Mean and Variance of the Solution

Let us consider the class of stochastic differential equations given by

$$dC(t) = [f(t) - \eta C(t)]\, dt + \sigma C(t)\, dW(t), \quad C(0) = c_0 > 0. \quad \text{(S21)}$$

**Mean:** Let $m(t) = \mathbb{E}[C(t)]$. Then $m(t)$ satisfies the ordinary differential equation given by

$$\frac{dm(t)}{dt} = f(t) - \eta m(t), \quad m(0) = c_0.$$

This is a first-order linear ordinary differential equation. Thus, the mean of Eq (S21) can be explicitly written as

$$m(t) = c_0 e^{-\eta t} + \int_0^t f(s)\, e^{-\eta(t-s)} ds. \quad \text{(S22)}$$

**Variance:** Consider $h(x) = x^2$. Using the Itô formulae on $h(C)$, we have

$$\begin{aligned} dh(C) &= h'(C)dC + \frac{1}{2}h''(C)(dC)^2 \\ &= \left[2Cf(t) - 2\eta C^2 + \sigma^2 C^2\right] dt + 2\sigma C^2 dW(t) \end{aligned}$$

Taking the expectation of both sides and using the fact that the Itô integral is zero mean, we have that

$$\frac{d(\mathbb{E}\left[C(t)^2\right])}{dt} = 2\mathbb{E}[C(t)]\, f(t) - 2\eta\mathbb{E}\left[C(t)^2\right] + \sigma^2\mathbb{E}\left[C(t)^2\right]$$

Let $v(t) = \mathbb{E}\left[C(t)^2\right]$ then the variance of the stochastic differential equation given by Eq (S21) solves the ODE given by

$$\frac{dv(t)}{dt} = 2m(t)f(t) + (\sigma^2 - 2\eta)v(t), \quad v_0 = c_0^2.$$

Noting that the above equation is a linear first order equation, we can use the integrating factor, $e^{-(\sigma^2-2\eta)t}$. Multiplying both sides of the above equation, we get

$$\begin{aligned} e^{-(\sigma^2-2\eta)t}\frac{dv(t)}{dt} - (\sigma^2 - 2\eta)e^{-(\sigma^2-2\eta)t}v(t) &= 2m(t)f(t)e^{-(\sigma^2-2\eta)t} \\ \frac{d}{dt}\left(e^{-(\sigma^2-2\eta)t}v(t)\right) &= 2m(t)f(t)e^{-(\sigma^2-2\eta)t} \end{aligned}$$

Thus,

$$\begin{aligned} v(t) &= c_0^2 e^{(\sigma^2-2\eta)t} + \int_0^t 2e^{(\sigma^2-2\eta)(t-s)} m(s)f(s)ds \\ &= c_0^2 e^{(\sigma^2-2\eta)t} + \int_0^t 2e^{(\sigma^2-2\eta)(t-s)} \left(c_0 e^{-\eta s} + \int_0^s f(u)\, e^{-\eta(s-u)} du\right) f(s)ds \end{aligned}$$

Thus, variance can be found to be

$$\begin{aligned}
Var(C(t)) &= v(t) - m(t)^2 \\
&= c_0^2 e^{(\sigma^2-2\eta)t} + \int_0^t 2e^{(\sigma^2-2\eta)(t-s)} m(s) f(s) ds - m(t)^2 \\
&= c_0^2 e^{(\sigma^2-2\eta)t} + \int_0^t 2e^{(\sigma^2-2\eta)(t-s)} \left( c_0 e^{-\eta s} + \int_0^s f(u)\, e^{-\eta(s-u)} du \right) f(s) ds \\
&\quad - \left( c_0 e^{-\eta t} + \int_0^t f(s)\, e^{-\eta(t-s)} ds \right)^2
\end{aligned}$$

#### S5.2 Hitting Time Analysis

##### S5.2.1 Mean Detection time

Let $\tau = \inf\{t > 0 \,|\, C(t) \geq 1\}$ be a stopping time. This represents the expected time for the biomarker levels to reach detection levels given underlying tumor dynamics. Here we assume that the underlying tumor population has reached the equilibrium state. As such, given the stationary state in the ODE, we have $f(t)$ would not have a time dependency and can be viewed as a constant, say $f^*$. In this sense, we can consider $C(0)$ as the mean biomarker proportion as the ODE reaches "close" to the equilibrium state. Eq. (S22) provides one such interpretation. Namely, at $t$ large enough for the ODE to reach equilibrium and with $\hat{c}_0$ referring to the proportion of biomarker at the very start of the ODE initiation,

$$\begin{aligned}
m(t) &= \hat{c}_0 e^{-\eta t} + f^* \int_0^t e^{-\eta(t-s)} ds \\
&= \hat{c}_0 e^{-\eta t} + \frac{f^*}{\eta} \left(1 - e^{-\eta t}\right) \\
&\to \frac{f^*}{\eta}.
\end{aligned}$$

We want to find $\mathbb{E}\left[\tau \,|\, C(0)\right]$ for our process. Consider a differentiable function $k(x)$ that is to be defined later. Via the Itô lemma,

$$\begin{aligned}
dk(C) &= k'(C) dC + \frac{1}{2} k''(C) (dC)^2 \\
&= \left( f^* k'(C) - \eta C k'(C) + \frac{1}{2} k''(C) \sigma^2 C^2 \right) dt + \sigma C k'(C) dB(t).
\end{aligned}$$

We integrate this SDE from 0 to $\tau$ to get

$$k(C) - k(c_0) = \int_0^\tau \left( f^* k'(C) - \eta C k'(C) + \frac{1}{2} k''(C) \sigma^2 C^2 \right) dt + \int_0^\tau \sigma C k'(C) dB(t).$$

Taking expectations on both sides and via the Optional Sampling Theorem for sub-martingales, we have

$$k(1) - k(c_0) = \mathbb{E}\left[ \int_0^\tau \left( f^* k'(C) - \eta C k'(C) + \frac{1}{2} k''(C) \sigma^2 C^2 \right) dt \right]$$

Thus, suppose that $k$ satisfies the ODE given by

$$\begin{cases} (f^* - \eta x) k'(x) + \dfrac{1}{2} k''(x) \sigma^2 x^2 = -1, \\ \qquad k(1) = 0, \quad k'(1) = 0. \end{cases} \tag{S23}$$

Then we can use this function and the initial population ratio to obtain the average time for detection. Namely,

$$\mathbb{E}\left[\tau \mid C(0)\right] = k(c_0).$$

Thus, we pivot to solving the differential equation presented above. We can rewrite Eq (S23) as

$$\begin{cases} k''(x) + p(x)k'(x) = q(x), \\ \quad k(1) = 0, \quad k'(1) = 0. \end{cases} \tag{S24}$$

where

$$p(x) = \frac{2(f^* - \eta x)}{\sigma^2 x^2}, \qquad q(x) = -\frac{2}{\sigma^2 x^2}. \tag{S25}$$

Note that both $p$ and $q$ are continuous when $x > 0$ and as such, there exists a unique solution to the second order non-homogeneous linear differential equation given by Eq. (S24). We start from the homogeneous form to obtain the general solution.

$$\begin{aligned} k'' + p(x)k' &= 0 \\ e^{-\int_x^1 p(v)dv}(k'' + p(x)k') &= 0 \\ e^{-\int_x^1 p(v)dv}k'' + p(x)e^{-\int_x^1 p(v)dv}k' &= 0 \\ (e^{-\int_x^1 p(v)dv}k')' &= 0 \\ k' &= -c_2 e^{\int_x^1 p(v)dv} \\ k(x) &= c_2 \int_x^1 e^{\int_y^1 p(v)dv} dy + c_1 \end{aligned}$$

where we have integrated from some arbitrary initial value $x \in (0, 1)$ to the final value $C(\tau) = 1$. Thus, we have the two homogeneous solutions given by

$$k_1(x) = 1, \qquad k_2(x) = \int_x^1 e^{\int_y^1 p(v)dv} dy.$$

We calculate the Wronskian for these two solutions:

$$\begin{aligned} W(x) &= k_1 k_2' - k_1' k_2 \\ &= k_2' \\ &= -e^{\int_x^1 p(v)dv} \\ &\neq 0 \, \forall \, x \in [0, 1] \end{aligned}$$

This means that the two solutions are fundamental solutions for the general case. We now use these general case solutions and method of variation of parameters to obtain the particular solution for the non-homogeneous case. It is well known that the particular solution will have the form

$$\begin{aligned} k_p(x) &= -k_1 \int_x^1 \frac{k_2(v)q(v)}{W(v)} dv + k_2 \int_x^1 \frac{k_1(v)q(v)}{W(v)} dv \\ &= -\int_x^1 \frac{k_2(v)q(v)}{W(v)} dv + k_2(x) \int_x^1 \frac{q(v)}{W(v)} dv \end{aligned} \tag{S26}$$

For the first term in the right hand side of the above equation,

$$
\begin{aligned}
-\int_x^1 \frac{k_2(v)q(v)}{W(v)}dv &= -\int_x^1 \frac{\int_v^1 e^{\int_y^1 p(z)dz}dy}{-e^{\int_v^1 p(z)dz}}\, q(v)dv \\
&= \int_x^1 \int_v^1 \frac{e^{\int_y^1 p(z)dz}}{e^{\int_v^1 p(z)dz}}dy\, q(v)dv \\
&= \int_x^1 \int_v^1 e^{-\int_v^y p(z)dz}dy\, q(v)dv \qquad \text{(S27)}
\end{aligned}
$$

Similarly from (S26),

$$
\begin{aligned}
\int_x^1 \frac{k_2(x)q(v)}{W(v)}dv &= \int_x^1 \frac{\int_x^1 e^{\int_y^1 p(z)dz}}{-e^{\int_v^1 p(z)dz}}dy\, q(v)dv \\
&= -\int_x^1 \int_x^1 e^{\int_y^1 p(z)dz - \int_v^1 p(z)dz}dy\, q(v)dv \\
&= -\int_x^1 \int_x^1 e^{-\int_v^y p(z)dz}dy\, q(v)dv \qquad \text{(S28)}
\end{aligned}
$$

Combining Eqs. (S26),(S27) and, (S28), we get

$$
\begin{aligned}
k_p(x) &= \int_x^1 \int_v^1 e^{-\int_v^y p(z)dz}dy\, q(v)dv - \int_x^1 \int_x^1 e^{-\int_v^y p(z)dz}dy\, q(v)dv \\
&= \int_x^1 \left[\int_v^1 e^{-\int_v^y p(z)dz}dy - \int_x^1 e^{-\int_v^y p(z)dz}dy\right] q(v)dv \\
&= -\int_x^1 \int_x^v e^{-\int_v^y p(z)dz}dy\, q(v)dv.
\end{aligned}
$$

Thus, the general form for our solution is given by

$$
\begin{aligned}
k(x) &= c_1k_1 + c_2k_2 + k_p \qquad \text{(S29)} \\
&= c_1 + c_2\int_x^1 e^{\int_y^1 p(v)dv}dy - \int_x^1 \int_x^v e^{-\int_v^y p(z)dz}dy\, q(v)dv, \qquad \text{(S30)}
\end{aligned}
$$

where $c_1$ and $c_2$ depend on the boundary conditions for $k(1)$ and $k'(1)$. From Eq. (S30) we can see that, $k(1) = 0$ implies that $c_1 = 0$. Furthermore, using FTC and Leibniz Integral Rule we get

$$
k'(x) = -c_2 e^{\int_x^1 p(v)dv} + \int_x^1 e^{-\int_v^x p(z)dz}q(v)dv + \int_x^1 e^{-\int_x^y p(z)dz}q(x)dy.
$$

As such, $k'(1) = 0$ implies that $c_2 = 0$. Thus, conditional on the fact that $C_0 = c_0 \in (0, 1]$, we can then find the mean time for escape to be given by

$$
\mathbb{E}\left[\tau \,|\, C(0) = c_0\right] = -\int_{c_0}^1 \int_{c_0}^v e^{-\int_v^y p(z)dz}dy\, q(v)dv
$$

where $p$, $q$ are as defined in Eq (S25).

##### S5.2.2 Mean "Extinction" Time and distribution

The solution of the stochastic differential equation defined by (S4) is given by Eq. (S20). As such, we can see that this process does not reach zero at all. However, consider $0 < \epsilon << 1$ and the associated stopping time $\tau_\epsilon = \inf\{t \geq 0 \,|\, C(t) \leq \epsilon\}$. Then we can turn our focus to the first time the stochastic process is close to zero.

**Hitting Probabilities:**

Using the exposition from Tan in [16], let $u_a(x,t)$ be the probability that the diffusion process $C(t)$ starting at $x > a$ hits $a$ before or at time $t$. More formally, we define the following transformation of the conditional pdf of $C(t)$,

$$u_a(x,t) = \int_0^\infty g(x,z;s,t)\delta(z-a)dz = g(x,a;s,t)$$

where $g$ is the conditional pdf of the diffusion process. In our case, we are looking for the case where $a = \epsilon$. From Theorem 6.2 and Section 7.3.1. of [16], we have that $u_\epsilon(x,t)$ satisfies the Kolmogorov backward equation. Namely,

$$\frac{\partial}{\partial t}u_\epsilon(x,t) = (f(t) - \eta x)\frac{\partial}{\partial x}u_\epsilon(x,t) + \frac{1}{2}\sigma^2 x^2 \frac{\partial^2}{\partial x^2}u_\epsilon(x,t) \tag{S31}$$

We define $U_\epsilon(p) = \lim_{t\to\infty} u_\epsilon(p,t)$ as the ultimate probability of elimination. Taking the limit as $t$ grows large on both sides of Eq. (S31), we first assume that $f(t)$ has already reached its limiting behavior $f^*$ and as such, we get

$$(f^* - \eta x)\frac{d}{dp}U_\epsilon(x) + \frac{1}{2}\sigma^2 x^2 \frac{d^2}{dx^2}U_\epsilon(x) = 0 \tag{S32}$$

with $U_\epsilon(0) = 0$ and $\lim_{x\to\infty} U_\epsilon(x) = 0$ as functions in $C[0,T]$. Using an integrating factor, it can be shown that the the solution of (S32) is given by

$$U_\epsilon(x) = 1 - \frac{\int_\epsilon^x \phi(y)dy}{\lim_{b\to\infty}\int_\epsilon^b \phi(y)dy}, \tag{S33}$$

where $\phi$ is given by

$$\begin{aligned}\phi(x) &= \exp\left\{-2\int_\epsilon^x \frac{f^* - \eta y}{\sigma^2 y^2}dy\right\},\\ &= \left(\frac{x}{\epsilon}\right)^{\frac{2\eta}{\sigma^2}} \times e^{-\frac{2f^*}{\sigma^2\epsilon}} \times e^{\frac{2f^*}{\sigma^2 x}}.\end{aligned} \tag{S34}$$

Note that $\phi$ is increasing for $x > \frac{f^*}{\eta}$ and as such, the denominator in Eq. (S33) grows to infinity leading to divergence in the limit. Thus,

$$U_\epsilon(x) = 1. \tag{S35}$$

**Hitting Times:**

We can exploit the simple nature of our SDE for some classical results about hitting times. From Section 5.5C in Karatzas and Shreve [17], for a diffusion process given by a general linear stochastic differential equation, we gave the mean hitting time is given by

$$\mathbb{E}\left[\tau_\epsilon | x\right] = -\int_\epsilon^x (\psi(x) - \psi(y))\,\mu(dy) + \lim_{b\to\infty}\frac{\psi(x)-\psi(\epsilon)}{\psi(b)-\psi(\epsilon)}\int_\epsilon^b (\psi(b)-\psi(y))\,\mu(dy)$$

where $\psi$ is the scale function defined as

$$\psi(x) = \int_{\epsilon}^{x} \phi(y)\,dy$$

with $\phi$ from Eq.(S34) and the speed measure, $\mu$, is given by

$$\begin{aligned}\mu(dx) &= \frac{2\,dx}{\psi'(x)\sigma^2 x^2}, \\ &= \frac{2\,dx}{\sigma^2 x^2 \phi(x)},\end{aligned}$$

as,

$$\psi'(x) = \phi(x).$$

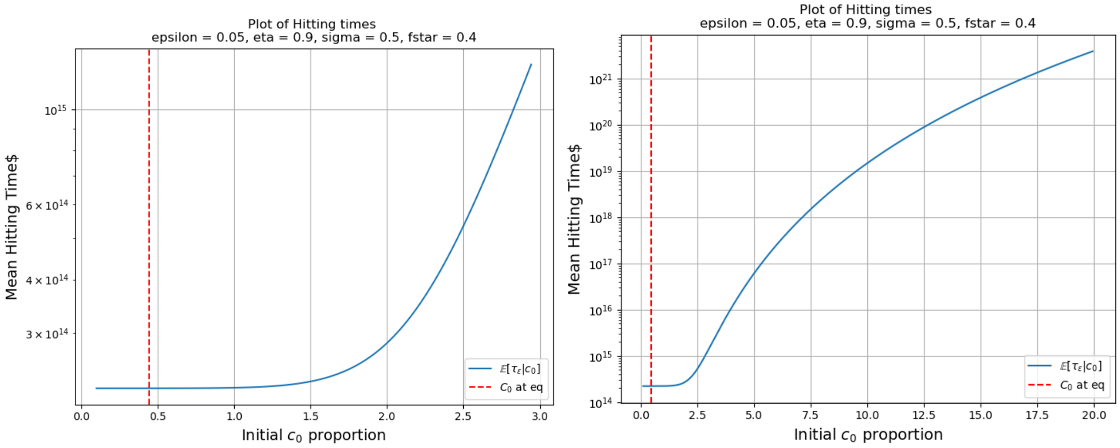


Figure S8: Mean Hitting Times for given params. The second panel is a zoomed out version of the first. These Blow up if epsilon is really small or $c_0$ large

#### S5.3 Anticipative Stochastic Differential Equation

Due to the lower detection limit for the biomarker, when we receive a signal, it represents biomarker activity leaving a certain size. In the previous section, we looked at the stopping time for when the ratio $c^*$ reaches 1. In this section, we flip the perspective to assume that the observed signal received at time $T$ is obtained at $t = 0$. Without loss of generality, we assume $T = 1$. Such perspective is used in financial mathematics to analyze insider trading. We use a similar motivation but for the biomarker compartment. As such, we try to look "forward" in time with "future" information with the help of the generalized Itô integral. To that extent, we apply apply anticipative stochastic calculus via the Ayed-Kuo stochastic integral [18]. We provide a gentle introduction below.

Consider $\{W_t, t \in [0,1]\}$, a fixed Brownian motion and the following $\sigma$-fields:

$$\mathcal{F}_s = \sigma\{W_u;\ 0 \le u \le s\}, \qquad 0 \le s \le 1$$
$$\mathcal{G}^{(t)} = \sigma\{W_1 - W_v;\ t \le v \le 1\}, \qquad 0 \le t \le 1$$

We call $\{\mathcal{F}_s : s \in [0,1]\}$ the *forward*-filtration and $\{\mathcal{G}^{(t)} : t \in [0,1]\}$ the *backward* or *counter*-filtration generated by the Brownian motion. Before we define the Ayed–Kuo integral, we need to define instantly independent processes. A stochastic process $\phi(t)$ is called *instantly independent* with respect to the forward filtration $\{\mathcal{F}_t\}$ if for each $t \in [0,1]$, the random variable $\phi(t)$ and the $\sigma$-field $\mathcal{F}_t$ are independent. We concisely state the key steps in the definition of the integral. For a more detailed description, please refer to [18, Section 2].

**Definition S5.1** ( [18, Definition 2.3]). *The anticipating integral is defined in following three steps:*

1. *Suppose $f(t)$ is an $\{\mathcal{F}_t\}$-adapted continuous stochastic process and $\phi(t)$ is a continuous stochastic processes that is instantly independent with respect to $\{\mathcal{F}_t\}$. Then the stochastic integral of $\Phi(t) = f(t)\phi(t)$ is defined by*

$$\int_0^1 f(t)\ \phi(t)\ dW_t = \lim_{\|\Delta_n\| \to 0} \sum_{j=1}^{n} f(t_{j-1})\ \phi(t_j)\ (W_{t_j} - W_{t_{j-1}}),$$

   *provided that the limit exists in probability.*

2. *For any stochastic process of the form $\Phi(t) = \sum_{i=1}^n f_i(t)\phi_i(t)$, where $f_i(t)$ and $\phi_i(t)$ are given as in step (1), the stochastic integral is defined by*

$$\int_0^1 \Phi(t)\ dW_t = \sum_{i=1}^{n} \int_0^1 f_i(t)\ \phi_i(t)\ dW_t.$$

3. *Let $\Phi(t)$ be a stochastic process such that there is a sequence $(\Phi_n(t))_{n=1}^{\infty}$ of stochastic processes of the form in step (2) satisfying*

   *(a)* $\int_0^1 |\Phi_n(t) - \Phi(t)|^2\, dt \to 0$ *almost surely as* $n \to \infty$, *and*

   *(b)* $\int_0^1 \Phi_n(t)\ dW_t$ *converges in probability as* $n \to \infty$.

   *Then the stochastic integral of $\Phi(t)$ is defined by*

$$\int_0^1 \Phi(t)\ dW_t = \lim_{n\to\infty} \int_0^1 \Phi_n(t)\ dW_t \quad \textit{in probability.}$$

In addition. we state an extension of Itô's formula that can account for instantly independent processes. Let $X_t$ and $Y^{(t)}$ be stochastic processes of the form

$$X_t = X_0 + \int_0^t g(s)\, dW_s + \int_0^t h(s)\, ds, \tag{S36}$$

$$Y^{(t)} = Y^{(1)} + \int_t^1 \xi(s)\, dW_s + \int_t^1 \eta(s)\, ds, \tag{S37}$$

where $g(t), h(t)$ are adapted (so $X_t$ is an Itô process), and $\xi(t), \eta(t)$ are instantly independent such that $Y^{(t)}$ is also instantly independent.

**Theorem S5.2** ( [18, Theorem 3.2]). *Suppose $\{X_t^{(i)}\}_{i=1}^n$ and $\{Y_j^{(t)}\}_{j=1}^m$ are of the form* (S36) *and* (S37)*, respectively. Suppose $\theta(t, x_1, \ldots, x_n, y_1, \ldots, y_m)$ is a real-valued function that is $C^1$ in $t$ and $C^2$ in other variables. Then the stochastic differential of $\theta(t, X_t^{(1)}, \ldots, X_t^{(n)}, Y_1^{(t)}, \ldots, Y_m^{(t)})$ is given by*

$$\begin{aligned} &d\theta(t, X_t^{(1)}, \ldots, X_t^{(n)}, Y_1^{(t)}, \ldots, Y_m^{(t)}) \\ &= \theta_t\, dt + \sum_{i=1}^n \theta_{x_i} dX_t^{(i)} + \sum_{j=1}^m \theta_{y_j} dY_j^{(t)} \\ &\quad + \frac{1}{2} \sum_{i,k=1}^n \theta_{x_i x_k} dX_t^{(i)} dX_t^{(k)} - \frac{1}{2} \sum_{j,l=1}^m \theta_{y_j y_l} dY_j^{(t)} dY_l^{(t)}. \end{aligned}$$

With this machinery in place, we now make a strong assumption that the initial condition is a continuous functional of the Brownian noise, $W_1$. As such, we have the following anticipative stochastic differential equation with anticipating initial condition;

$$\begin{cases} dC_t = [f(E, B, I) - \eta C]\, dt + \sigma C\, dW(t), \quad t \in [0, 1], \\ C_0 = \psi(\xi W_1), \end{cases} \tag{S38}$$

We start off by assuming the form of the solution and then apply the general Itô formula given in Theorem S5.2 to find a solution that solves the linear stochastic differential equation. Motivated by the non-anticipatory case given in Eq. S20 suppose

$$\begin{aligned} Z(t) = \psi(\xi W_1 - Q(t)) e^{\sigma\, W_t - (\eta + \frac{1}{2}\sigma^2)t} \\ + \int_0^t e^{\sigma\, (W_t - W(s)) - (\eta + \frac{1}{2}\sigma^2)(t-s)} f\,(s)\; ds. \end{aligned} \tag{S39}$$

We need to determine the Itô process $Q(t)$ with $Q(1) = 0$. In order to apply the generalized Itô formula, we write

$$\begin{aligned} Z(t) = \psi(\xi\, (W_1 - W_t) + \xi W_t - Q(t)) e^{\sigma\, W_t - (\eta + \frac{1}{2}\sigma^2)t} \\ + \int_0^t e^{\sigma\, (W_t - W(s)) - (\eta + \frac{1}{2}\sigma^2)(t-s)} f\,(s)\; ds \end{aligned} \tag{S40}$$

We define the instantly independent process $Y^{(t)} = \xi\, (W_1 - W_t)$ and the following adapted processes

$$\begin{aligned} X_t^{(1)} &= e^{\sigma\, W_t - (\eta + \frac{1}{2}\sigma^2)t}, \\ X_t^{(2)} &= \xi W_t - Q(t), \quad \text{and} \\ X_t^{(3)} &= \int_0^t e^{\sigma\, (W_t - W(s)) - (\eta + \frac{1}{2}\sigma^2)(t-s)} f\,(s)\; ds. \end{aligned}$$

From the definitions of $X_t^{(1)}$, $X_t^{(2)}$, and $Y^{(t)}$ above, we get the differentials

$$
\begin{aligned}
dX_t^{(1)} &= \sigma X_t^{(1)}\, dW_t - \eta X_t^{(1)}\, dt, \\
dX_t^{(2)} &= \xi\, dW_t - dQ(t), \\
dX_t^{(3)} &= \left(f(t) - \eta X_t^{(3)}\right) dt + \sigma X_t^{(3)} dW_t, \\
(dX_t^{(1)})^2 &= \sigma^2 (X_t^{(1)})^2\, dt, \\
(dX_t^{(2)})^2 &= \xi^2\, dt - 2\xi\, dW_t dQ(t) + (dQ(t))^2, \\
dX_t^{(1)} dX_t^{(2)} &= \sigma\xi X_t^{(1)}\, dt - \sigma X_t^{(1)}\, dW_t dQ(t) + \eta X_t^{(1)}\, dQ(t)dt, \\
dY^{(t)} &= -\xi\, dW_t, \\
(dY^{(t)})^2 &= \xi^2\, dt.
\end{aligned}
$$

Now, define $\theta(x_1, x_2, x_3, y) = \psi(x_2 + y)x_1 + x_3$, so that $Z(t) = \theta\left(X_t^{(1)}, X_t^{(2)}, X_t^{(3)}, Y^{(t)}\right)$. From this, we get the partial derivatives

$$
\begin{aligned}
\theta_{x_1} &= \psi, & \theta_{x_1 x_1} &= 0, \\
\theta_{x_2} &= \psi' x_1, & \theta_{x_2 x_2} &= \psi'' x_1, \\
\theta_{x_3} &= 1, & \theta_{x_3 x_3} &= 0, \\
\theta_y &= \psi' x_1, & \theta_{yy} &= \psi'' x_1 \\
\theta_{x_1 x_2} &= \psi', & \theta_{x_3 x_2} &= 0 = \theta_{x_3 x_1}.
\end{aligned}
$$

Applying Theorem S5.2 and putting everything together, we can easily find the stochastic differential of $Z(t)$:

$$
\begin{aligned}
dZ(t) &= d\theta(X_t^{(1)}, X_t^{(2)}, Y^{(t)}) \\
&= \theta_{x_1} dX_t^{(1)} + \theta_{x_2} dX_t^{(2)} + \theta_{x_3} dX_t^{(3)} \\
&\quad + \frac{1}{2}\theta_{x_1 x_1}(dX_t^{(1)})^2 + \frac{1}{2}\theta_{x_2 x_2}(dX_t^{(2)})^2 + \frac{1}{2}\theta_{x_3 x_3}(dX_t^{(3)})^2 \\
&\quad + \theta_{x_1 x_2}(dX_t^{(1)})(dX_t^{(2)}) + \theta_{x_1 x_3}(dX_t^{(1)})(dX_t^{(3)}) + \theta_{x_3 x_2}(dX_t^{(3)})(dX_t^{(2)}) \\
&\quad + \theta_y\, dY^{(t)} - \frac{1}{2}\theta_{yy}(dY^{(t)})^2 \\
&= \psi\left(\sigma X_t^{(1)}\, dW_t - \eta X_t^{(1)}\, dt\right) + \psi' \cdot X_t^{(1)}\, (\xi dW_t - dQ(t)) \\
&\quad + \left(f(t) - \eta X_t^{(3)}\right) dt + \sigma X_t^{(3)} dW_t + 0 \\
&\quad + \frac{1}{2}\psi'' \cdot X_t^{(1)} \left(\xi^2\, dt - 2\xi\, dW_t dQ(t) + (dQ(t))^2\right) + 0 \\
&\quad + \psi'\left(\sigma\xi X_t^{(1)}\, dt - \sigma X_t^{(1)}\, dW_t dQ(t) + \eta X_t^{(1)}\, dQ(t)dt\right) + 0 + 0 \\
&\quad - \xi\psi' \cdot X_t^{(1)}\, dW_t - \frac{1}{2}\xi^2\psi'' \cdot X_t^{(1)}\, dt \\
&= \sigma(\psi X_t^{(1)} + X_t^{(3)})\, dW_t + \left(f - \eta\left(\psi X_t^{(1)} + X_t^{(3)}\right)\right) dt \\
&\quad + \psi' X_t^{(1)} \left(\sigma\xi\, dt - \sigma\, dW_t dQ(t) + \eta\, dQ(t)dt - dQ(t)\right) \\
&\quad \frac{1}{2}\psi''\left(-2\xi\, dW_t dQ(t) + (dQ(t))^2\right)
\end{aligned} \tag{S41}
$$

Suppose,

$$dQ(t) = \sigma\xi\, dt - \sigma\, dW_t dQ(t) + \eta\, dQ(t)dt \tag{S42}$$

$$(dQ(t))^2 = 2\xi\, dW_t dQ(t) \tag{S43}$$

then, using the definition of $\theta$, S41 can be written as

$$Z(t) = \sigma Z(t)\, dW_t + (f - \eta Z(t))\; dt, \tag{S44}$$

which is nothing but Eq. S38. Thus, all that is left to do is find the form of the Itô term $Q(t)$ that satisfies Eq. S42 and Eq. S43. Firstly, let us assume that $dQ(t)$ has a Brownian derivative term. Then $dQ(t)\, dW_t = \gamma(t)\, dt$ for some $\gamma(t)$. Consequently, $dQ(t) = (\xi - \gamma(t))\sigma\, dt$, which contradicts the assumption that $Q(t)$ has a Brownian term. However, if $Q(t)$ is deterministic then,

$$dQ(t) = \sigma\xi\, dt$$

$$(dQ(t))^2 = 0 = 2\xi\, dW_t dQ(t)$$

Imposing the initial condition $Q(0) = 0$, we get that $Q(t) = \int_0^t \sigma\xi\, dt = \sigma\xi t$. Putting this in the assumed form of the solution, we get our result. Summarizing the above discussion, we have following theorem.

**Theorem S5.3.** *Given the following stochastic differential equation with anticipating initial condition,*

$$\begin{cases} dC_t = [f(E,B,I) - \eta C]\; dt + \sigma C\, dW(t), \quad t \in [0,1], \\ C_0 = \psi(\xi W_1), \end{cases} \tag{S45}$$

*where $f$ is defined as earlier and $\psi \in \mathbb{C}^2(\mathbb{R})$. Then, the solution in probability is given by*

$$C(t) = \psi(\xi W_1 - \sigma\xi t)e^{\sigma\, W_t - (\eta + \frac{1}{2}\sigma^2)t} + \int_0^t e^{\sigma\,(W_t - W(s)) - (\eta + \frac{1}{2}\sigma^2)(t-s)} f\,(s)\; ds, \quad t \in [0,1]. \tag{S46}$$

When we compare this result to the non-anticipating case in Eq. (S20), we see that the solution has a linear negative drift scaled by volatility from the initial anticipating conditions as well as the driving Brownian motion. This correction term removes the bias generated in using the future information. In Itô calculus, we model as if we were a blind walker: at time $t$, we only know the past and present. We can't see into the future. So when we integrate, we are only allowed to depend on what we know up to time $t$. However. in the Ayed-Kuo stochastic calculus, we are essentially an informed insider: at time $t$, we may already know something about the future at time 1. So now, when we integrate the function, we can "cheat" by using future information. This cheating creates bias in the integral — and the correction term precisely removes that bias to maintain consistency with stochastic calculus rules.

### S6 Alternate non-linear formulation of the SDE

Instead of the linear stochastic differential equation given in Eq. S4, we can instead consider

$$\begin{aligned} dC =& [f(E,B,I) - \eta C]\, dt + \sigma\sqrt{C} dW(t). \\ =& \eta\left[\frac{f}{\eta} - C\right] dt + \sigma\sqrt{C} dW(t). \end{aligned} \tag{S47}$$

Note that this is nothing but a general form of the Cox-Ingersoll-Ross (CIR) process where the constant $\alpha$ is replaced with a time dependent positive drift $\frac{f}{\eta}$. In quantitative finance, the CIR process is traditionally used to model the evolution of interest rates in a one factor model where the trajectory of the interest rate is driven by only one market risk [19, 20]. There is a rich history of research into the CIR process. The non-linear formulation for the diffusion term leads to an inability to obtain a closed from path-wise solution. However, one can obtain the exact transition distribution and generate exact samples for simple functional forms of $f$. In addition, the CIR process is non-negative if it satisfies the Feller condition,

$$2\eta\alpha = 2f > \sigma^2.$$

This condition can not be easily guaranteed for time-dependent $f$ through the whole trajectory of the stochastic process. As such, neither can non-negativity of the CIR process. Using the SDE given in Eq. (S47) we obtain the first and second moment formulas, which we show below.

#### S6.1 Mean and Variance

**Mean:** Let $m(t) = \mathbb{E}\left[C(t)\right]$ then $m(t)$ satisfies the ordinary differential equation given by

$$\frac{dm(t)}{dt} = f(t) - \eta m(t), \quad m_0 = c_0.$$

This is nothing but the same first order linear ordinary differential equation as in the case of the linear stochastic differential equation case. Thus, the mean of can be explicitly written as

$$m(t) = c_0 e^{-\eta t} + \int_0^t f(s)\, e^{-\eta(t-s)} ds. \tag{S48}$$

**Variance:** Consider $h(x) = x^2$. Using the Itô formulae on $h(C)$, we have

$$\begin{aligned} dh(C) &= h'(C)dC + \frac{1}{2}h''(C)(dC)^2 \\ &= \left[2Cf(t) - 2\eta C^2 + \sigma^2 C\right] dt + 2\sigma C dW(t) \end{aligned}$$

Taking the expectation of both sides and using the fact that the Itô integral is zero mean, we have that

$$\frac{d(\mathbb{E}\left[C(t)^2\right])}{dt} = 2\mathbb{E}\left[C(t)\right] f(t) - 2\eta \mathbb{E}\left[C(t)^2\right] + \sigma^2 \mathbb{E}\left[C(t)\right]$$

Let $v(t) = \mathbb{E}\left[C(t)^2\right]$ then the variance of the stochastic differential equation given by Eq (S48) solves the ODE given by

$$\frac{dv(t)}{dt} = \left(2f(t) + \sigma^2\right) m(t) - 2\eta v(t), \quad v_0 = c_0^2.$$

Noting that the above equation is a linear first order equation, we can use the integrating factor, $e^{2\eta t}$. Multiplying both sides of the above equation, we get

$$\begin{aligned} e^{2\eta t}\frac{dv(t)}{dt} + 2\eta e^{2\eta t} v(t) &= \left(2f(t) + \sigma^2\right) m(t) e^{2\eta t} \\ \frac{d}{dt}\left(e^{2\eta t} v(t)\right) &= \left(2f(t) + \sigma^2\right) m(t) e^{2\eta t} \end{aligned}$$

Thus,

$$\begin{aligned} v(t) &= c_0^2 e^{-2\eta t} + \int_0^t \left(2f(s) + \sigma^2\right) e^{-2\eta(t-s)} m(s) ds \\ &= c_0^2 e^{-2\eta t} + \int_0^t \left(2f(s) + \sigma^2\right) \left(c_0 e^{-\eta s} + \int_0^s f(u)\, e^{-\eta(s-u)} du\right) f(s) ds \end{aligned}$$

Thus, variance can be found to be

$$
\begin{aligned}
Var(C(t)) &= v(t) - m(t)^2 \\
=& c_0^2 e^{-2\eta t} + \int_0^t \left(2f(s) + \sigma^2\right) \left( c_0 e^{-\eta s} + \int_0^s f(u)\, e^{-\eta(s-u)} du \right) f(s) ds - m(t)^2 \\
=& c_0^2 e^{-2\eta t} + \int_0^t \left(2f(s) + \sigma^2\right) \left( c_0 e^{-\eta s} + \int_0^s f(u)\, e^{-\eta(s-u)} du \right) f(s) ds \\
&- \left( c_0 e^{-\eta t} + \int_0^t f(s)\, e^{-\eta(t-s)} ds \right)^2
\end{aligned}
$$

### S7 Figures and Results for Biomarker Profile ($\sigma^2 < 2\eta$)

- **Interior State:**

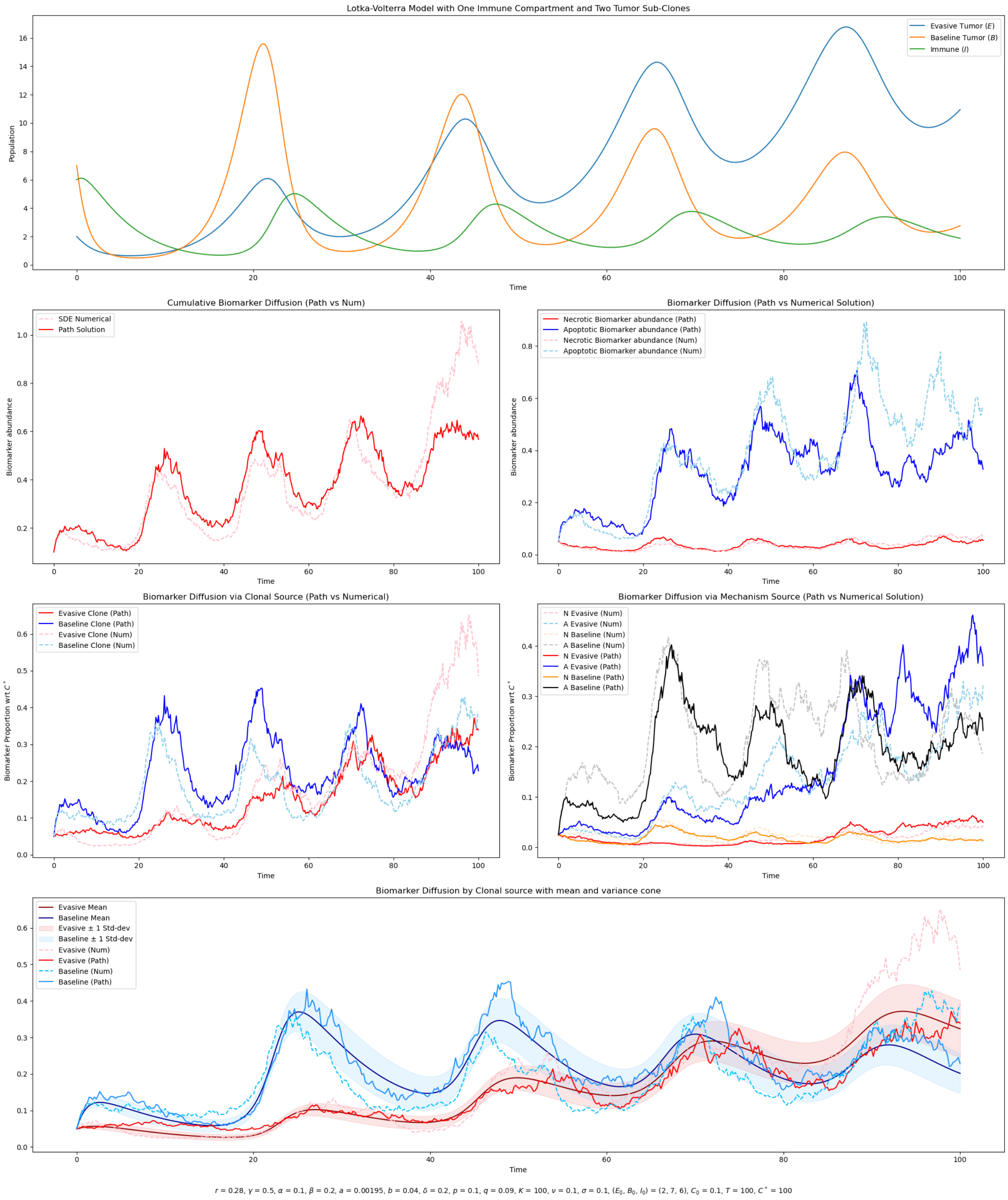


Figure S9: Biomarker Dynamics for Interior Equilibrium state. Parameter choices provided at the bottom of the figure

- **Evasive Loses, Baseline - Tumor Coexistence:**

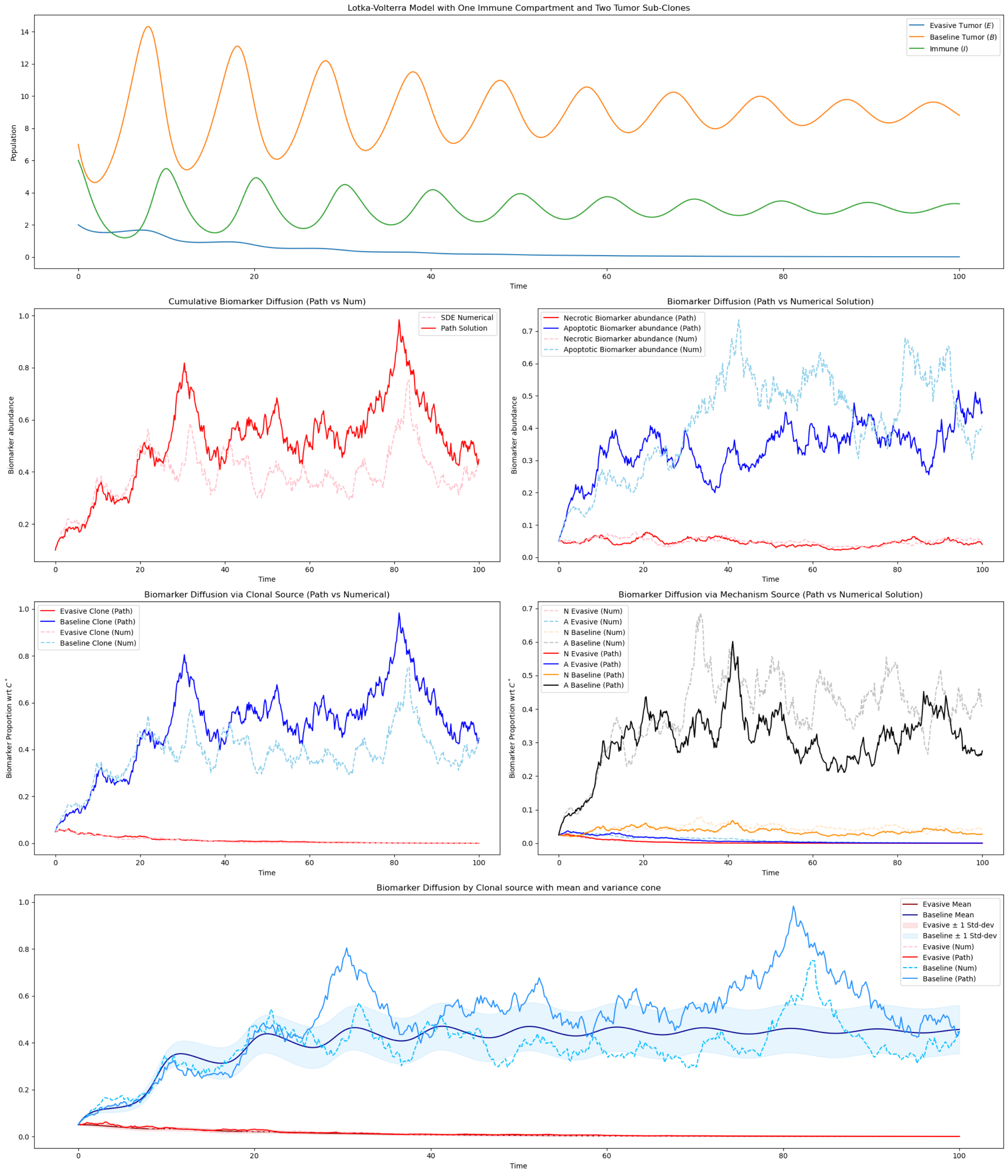


$r = 0.1$, $\gamma = 0.5$, $\alpha = 0.05$, $\beta = 0.15$, $a = 0.007$, $b = 0.1$, $\delta = 0.9$, $p = 0.5$, $q = 1.5$, $K = 100$, $\nu = 0.1$, $\sigma = 0.1$, $(E_0, B_0, I_0) = (2, 7, 6)$, $C_0 = 0.1$, $T = 100$, $C^* = 100$

Figure S10: Biomarker Dynamics for the case where evasive tumor Loses. Parameter choices provided at the bottom of the figure

- **Baseline loses, Evasive - Immune Coexistence:**

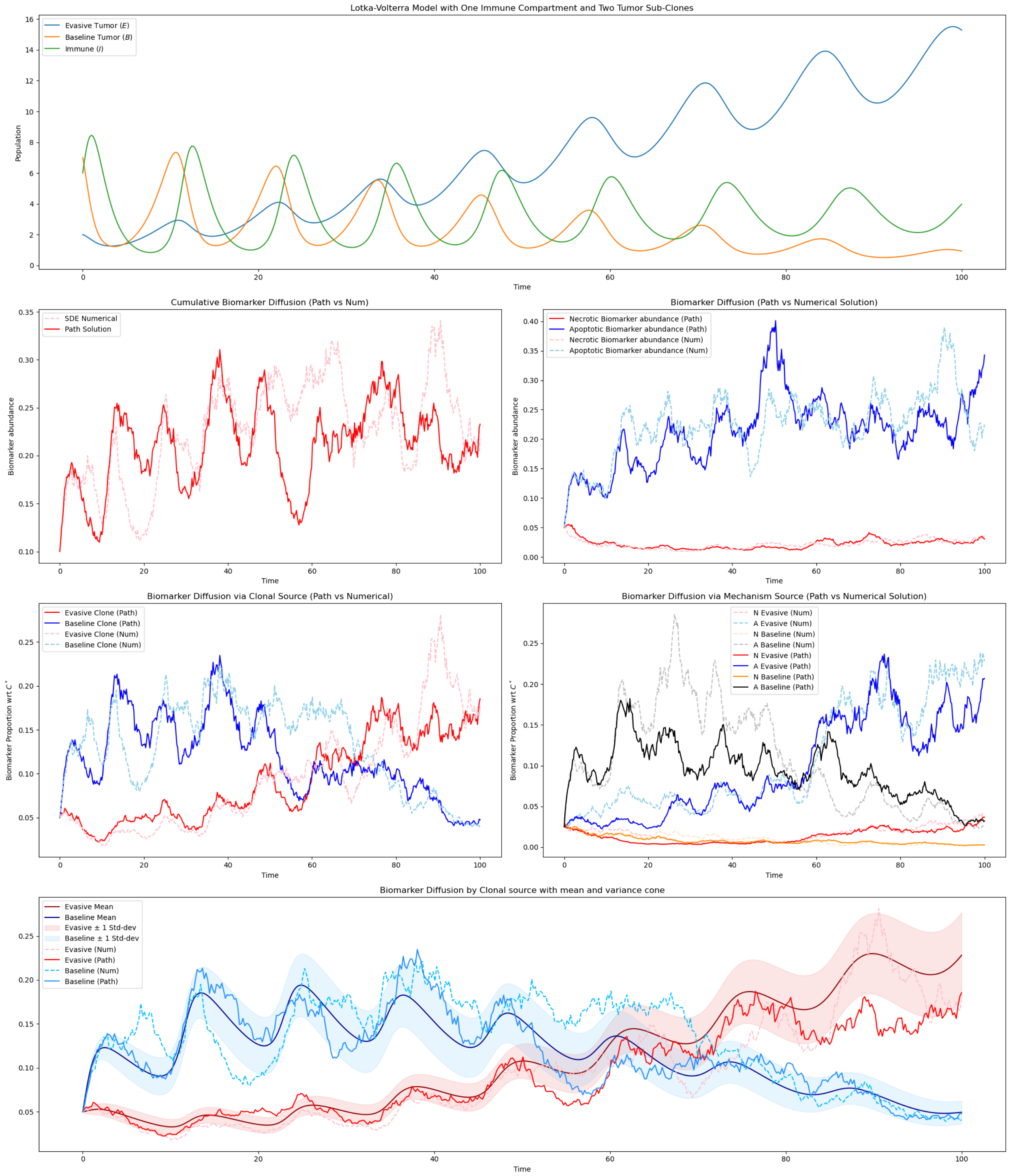


$r = 0.2$, $\gamma = 0.5$, $\alpha = 0.05$, $\beta = 0.15$, $a = 0.05$, $b = 0.2$, $\delta = 0.8$, $p = 0.4$, $q = 0.5$, $K = 100$, $\nu = 0.1$, $\sigma = 0.1$, $(E_0, B_0, I_0) = (2, 7, 6)$, $C_0 = 0.1$, $T = 100$, $C^* = 100$

Figure S11: Biomarker Dynamics for the case with evasive-immune coexistence. Parameter choices provided at the bottom of the figure

- **Immune loss:**

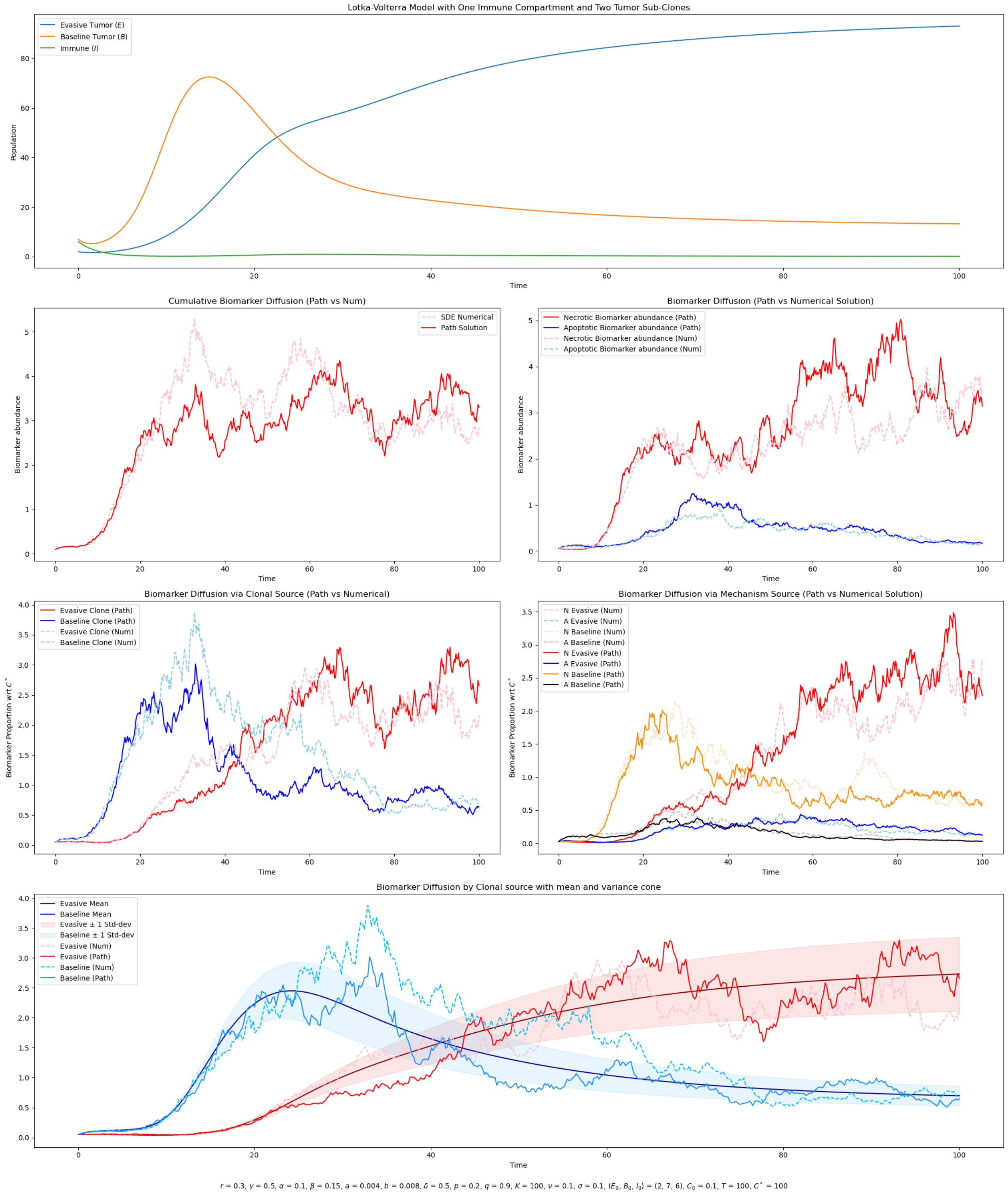


Figure S12: Biomarker Dynamics for the case where immune compartment loses. Parameter choices provided at the bottom of the figure

- **Evasive Win:**

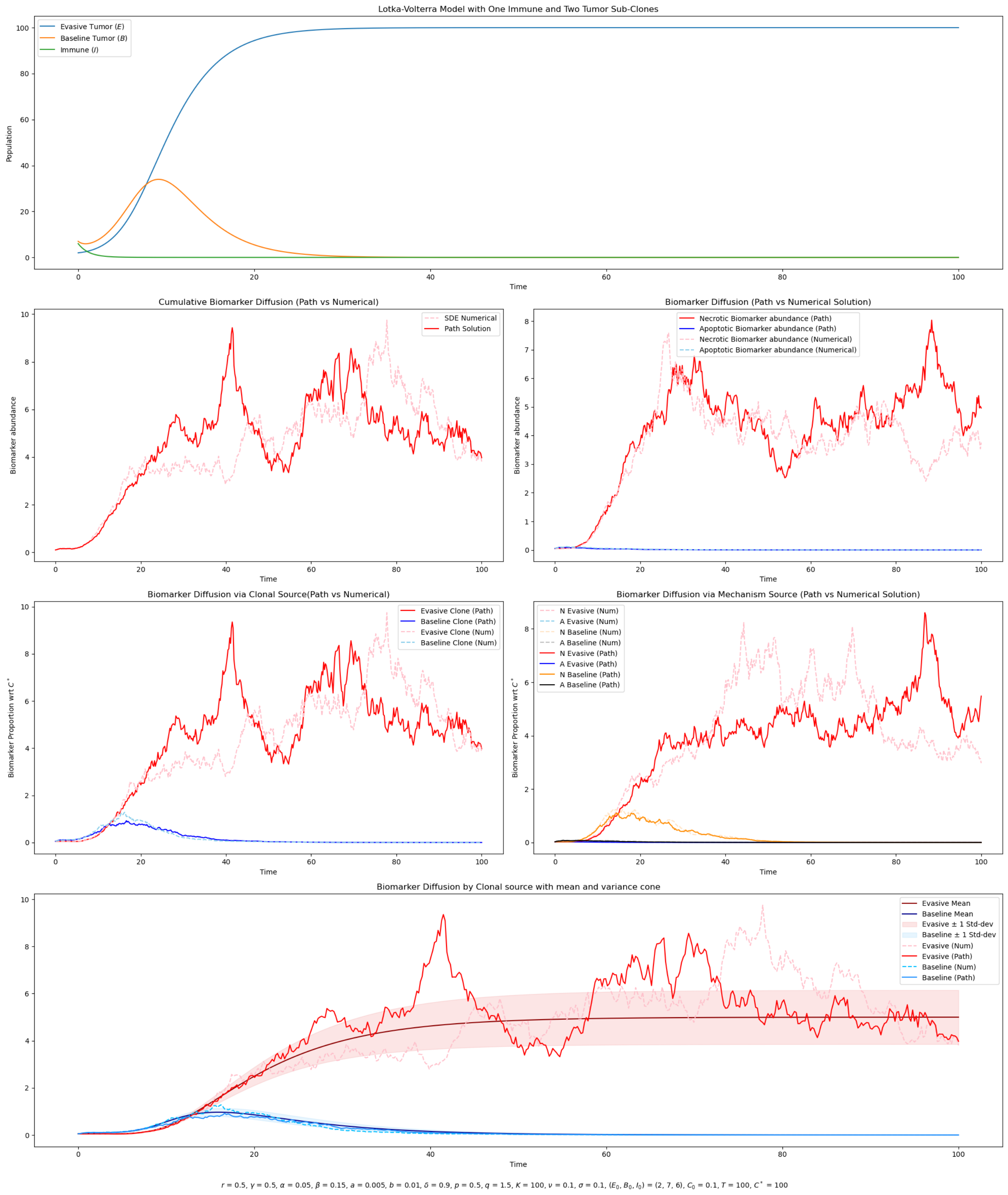


Figure S13: Biomarker Dynamics for the case with evasive tumor subpopulation win. Parameter choices provided at the bottom of the figure

- **Baseline Wins**

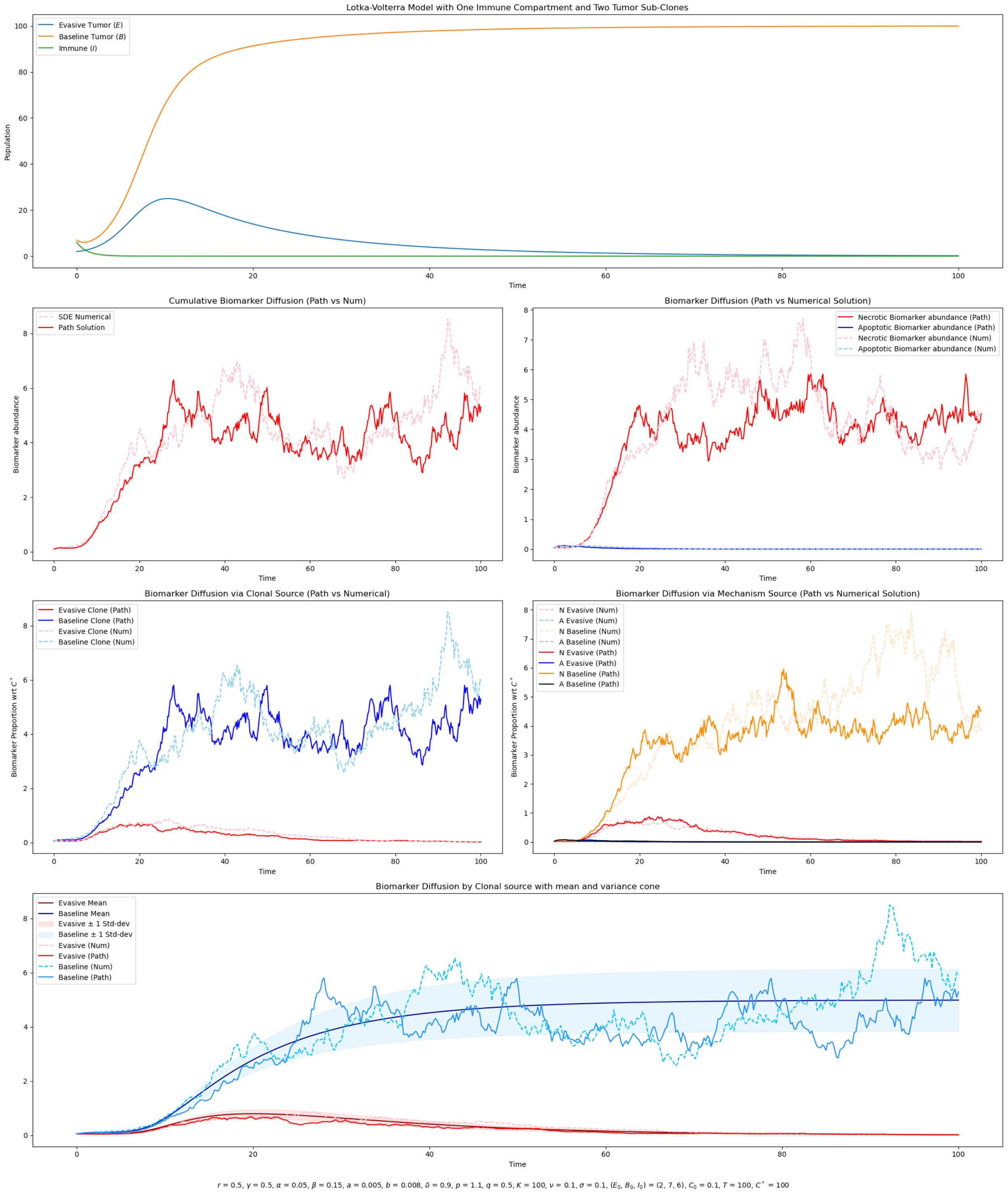


Figure S14: Biomarker Dynamics for the case where baseline wins. Parameter choices provided at the bottom of the figure

##### S7.0.1 Hitting Time behavior at Equilibria

- **Interior State:**

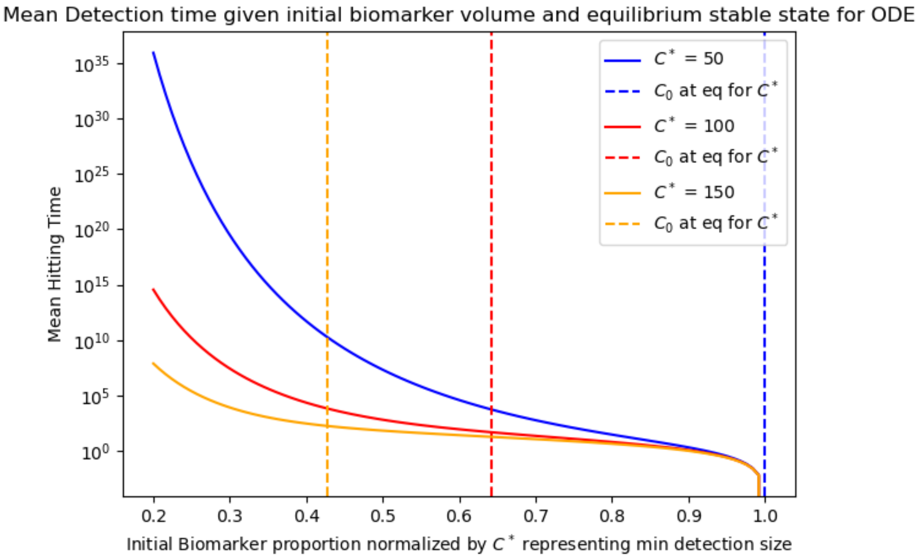


Figure S15: Interior State: Mean time to Detection after hitting interior equilibrium given initial $c_0$ values

- **Evasive Loses, Baseline-Tumor Coexistence:**

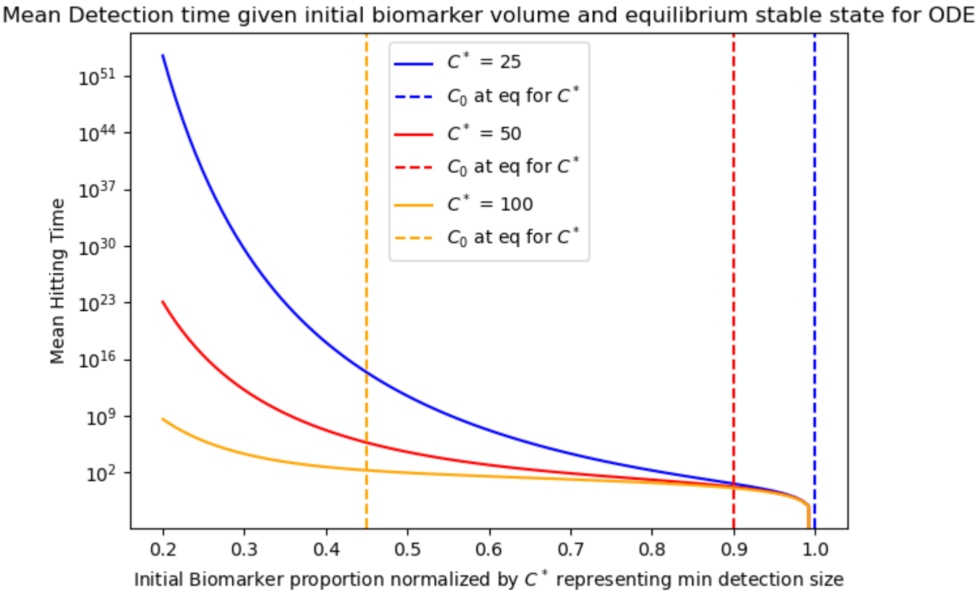


Figure S16: Baseline-Tumor Coexistence: Mean time to Detection after hitting equilibrium given initial $c_0$ values

- **Baseline loses, Evasive - Immune Coexistence:**

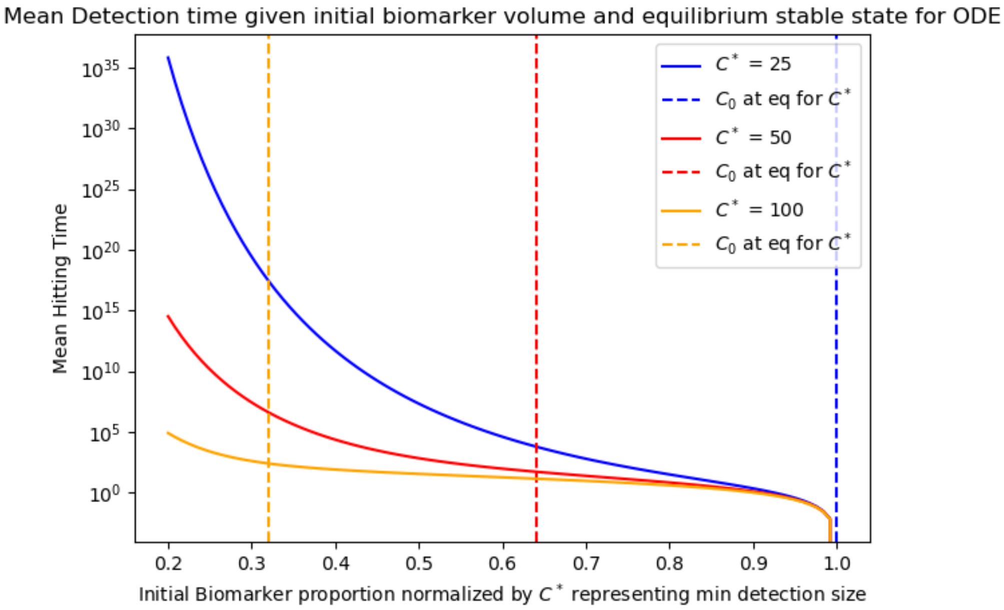


Figure S17: Evasive - Immune Coexistence: Mean time to Detection after hitting equilibrium given initial $c_0$ values

- **Immune loss:**

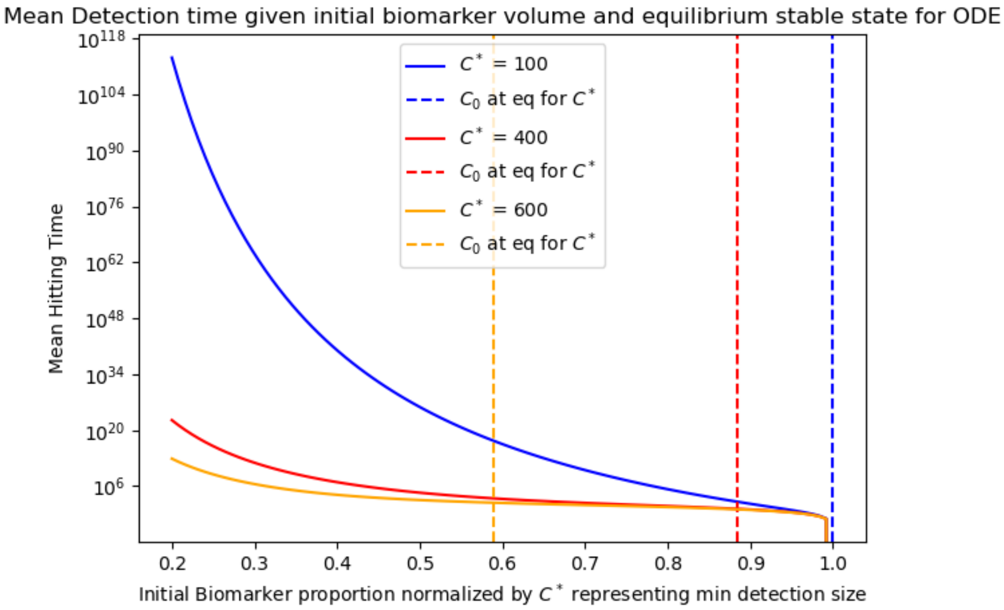


Figure S18: Immune loss: Mean time to Detection after hitting equilibrium given initial $c_0$ values

- **Evasive Win:**

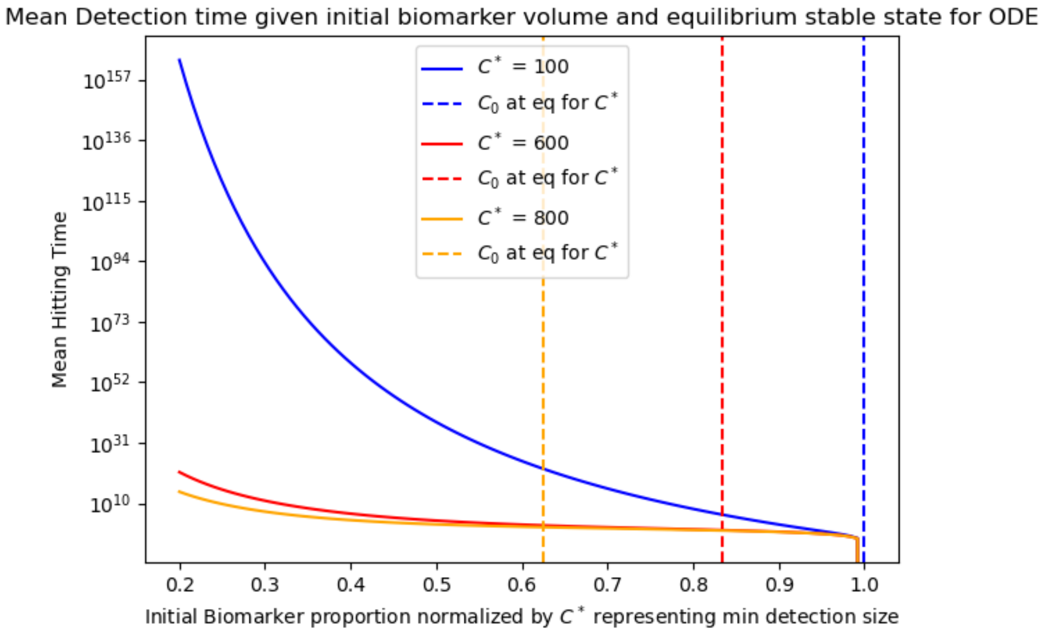


Figure S19: Evasive Win: Mean time to Detection after hitting equilibrium given initial $c_0$ values

- **Baseline Win:**

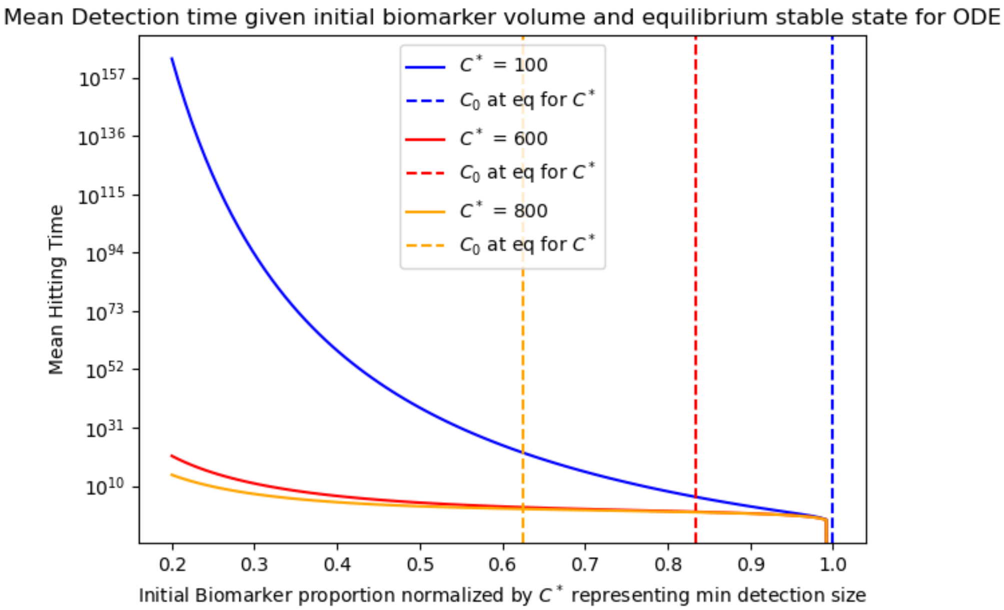


Figure S20: Baseline Win: Mean time to Detection after hitting equilibrium given initial $c_0$ values

##### S7.0.2 Effects on Volatility

In the previous simulations, we have that $\sigma^2 < 2\eta$ and as such, the decay rate of the biomarker mollifies the volatility as the time grows large. When the inequality is switched and the process is allowed to be more volatile as time grow, we find more earlier readouts. We plot the interior state case as an example,

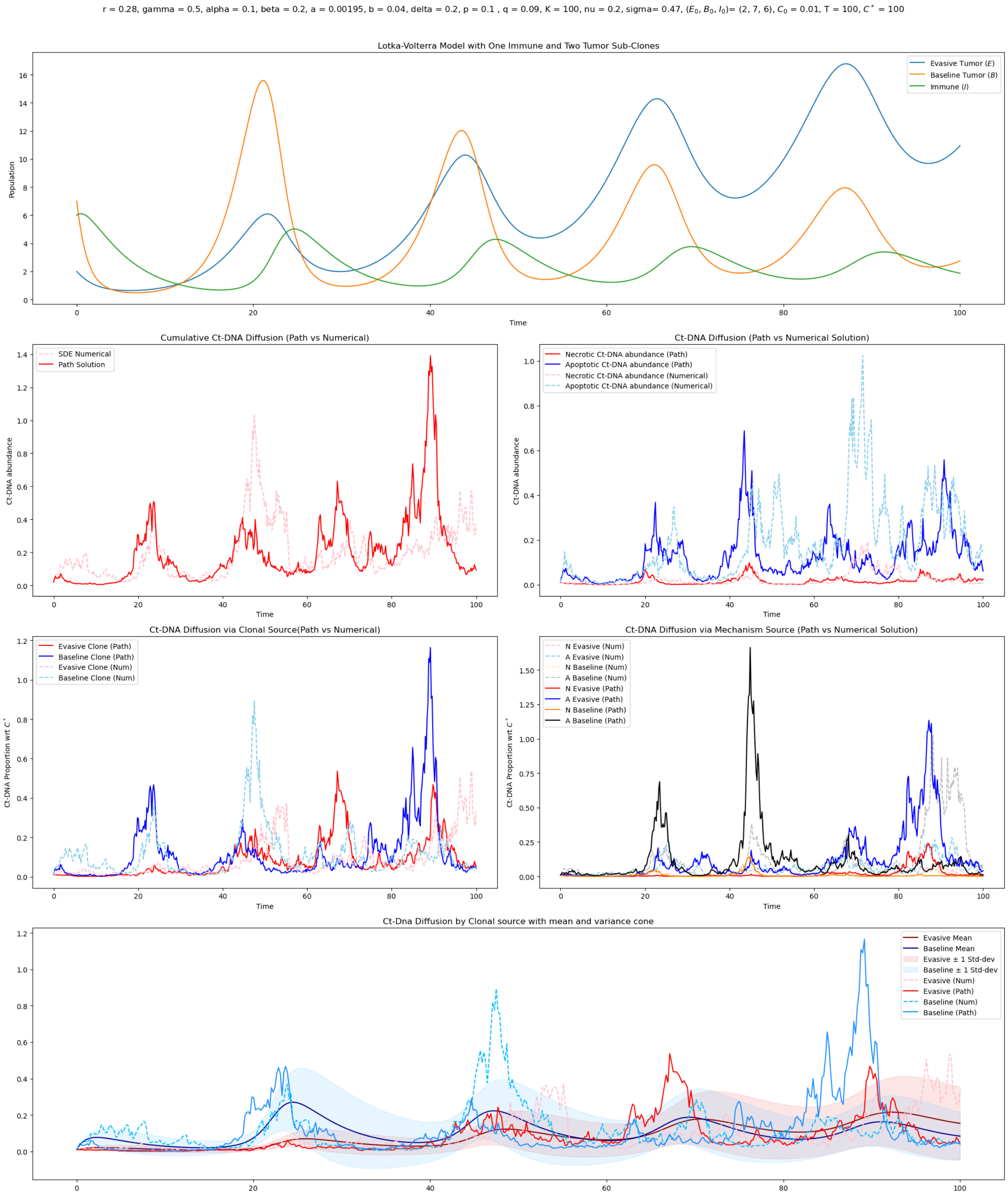


Figure S21: Biomarker Dynamics for the case with $\sigma^2 > 2\eta$. Parameters provided at the top of the figure.

### References


[1] Paras Jain, Ramanarayanan Kizhuttil, Madhav B Nair, Sugandha Bhatia, Erik W Thompson, Jason T George, and Mohit Kumar Jolly. Cell-state transitions and density-dependent interactions together explain the dynamics of spontaneous epithelial-mesenchymal heterogeneity. *iScience*, 2024.

[2] Muhammad Sufyan Bin Masroni, Kee Wah Lee, Victor Kwan Min Lee, Siok Bian Ng, Chao Teng Law, Kok Siong Poon, Bernett Teck-Kwong Lee, Zhehao Liu, Yuen Peng Tan, Wee Ling Chng, et al. Dynamic altruistic cooperation within breast tumors. *Molecular Cancer*, 22(1):206, 2023.

[3] Jordan Goldstein, Won Seog Kim, Sang Eun Yoon, Seok Jin Kim, Mark Roschewski, Jason Westin, Ryan C Lynch, Stefan K Alig, Sandra Close, Jacob J Chabon, et al. Optimizing circulating tumor dna limits of detection for dlbcl during first line therapy. *Blood*, 142:187, 2023.

[4] Pavel Stejskal, Hani Goodarzi, Josef Srovnal, Marián Hajdúch, Laura J van't Veer, and Mark Jesus M Magbanua. Circulating tumor nucleic acids: biology, release mechanisms, and clinical relevance. *Molecular cancer*, 22(1):15, 2023.

[5] Stefano Avanzini, David M Kurtz, Jacob J Chabon, Everett J Moding, Sharon Seiko Hori, Sanjiv Sam Gambhir, Ash A Alizadeh, Maximilian Diehn, and Johannes G Reiter. A mathematical model of ctDNA shedding predicts tumor detection size. *Sci Adv*, 6(50), December 2020.

[6] Aaron Li, Emil Lou, Kevin Leder, and Jasmine Foo. Early ctdna kinetics as a dynamic biomarker of cancer treatment response. *bioRxiv*, 2024.

[7] Thomas Rachman, David Bartlett, William LaFramboise, Patrick Wagner, Russell Schwartz, and Oana Carja. Modeling the effect of spatial structure on solid tumor evolution and circulating tumor dna composition. *Cancers*, 16(5), 2024.

[8] Grazziela P Figueredo, Peer-Olaf Siebers, and Uwe Aickelin. Investigating mathematical models of immuno-interactions with early-stage cancer under an agent-based modelling perspective. *BMC bioinformatics*, 14(Suppl 6):S6, 2013.

[9] Denise Kirschner and John Carl Panetta. Modeling immunotherapy of the tumor–immune interaction. *Journal of mathematical biology*, 37:235–252, 1998.

[10] Robert A Gatenby and Edward T Gawlinski. A reaction-diffusion model of cancer invasion. *Cancer research*, 56(24):5745–5753, 1996.

[11] Kathleen Wilkie. A review of mathematical models of cancer-immune interactions in the context of tumor dormancy. *Advances in experimental medicine and biology*, 734:201–234, 01 2013.

[12] Grace E Mahlbacher, Kara C Reihmer, and Hermann B Frieboes. Mathematical modeling of tumor-immune cell interactions. *Journal of Theoretical Biology*, 469:47–60, 2019.

[13] L. Perko. *Differential Equations and Dynamical Systems*. Texts in Applied Mathematics. Springer New York, 2012.

[14] J. J. Anagnost and C. A. Desoer. An elementary proof of the routh-hurwitz stability criterion. *Circuits, Systems and Signal Processing*, 10(1):101–114, Mar 1991.

[15] H. H. Kuo. *Introduction to stochastic integration*. Springer Universitext, 2006.

[16] W.Y. Tan. *Stochastic Models with Applications to Genetics, Cancers, AIDS and Other Biomedical Systems (second Edition)*. Series on concrete and applicable mathematics. World Scientific Publishing Company Pte Limited, 2015.

[17] Ioannis Karatzas and Steven Shreve. *Brownian motion and stochastic calculus*, volume 113. springer, 2014.

[18] Chii-Ruey Hwang, Hui-Hsiung Kuo, Kimiaki Saitô, and Jiayu Zhai. A general itô formula for adapted and instantly independent stochastic processes. *Communications on Stochastic Analysis*, 10(3), 2016.

[19] John C Cox, Jonathan E Ingersoll, Stephen A Ross, et al. A theory of the term structure of interest rates. *Econometrica*, 53(2):385–407, 1985.

[20] Damiano Brigo and Fabio Mercurio. A deterministic–shift extension of analytically–tractable and time–homogeneous short–rate models. *Finance and Stochastics*, 5:369–387, 2001.
